# The secretome of irradiated peripheral mononuclear cells attenuates hypertrophic skin scarring

**DOI:** 10.1101/2022.12.01.518726

**Authors:** Vera Vorstandlechner, Dragan Copic, Katharina Klas, Martin Direder, Golabi, Christine Radtke, Hendrik J. Ankersmit, Michael Mildner

**Affiliations:** Laboratory for Cardiac and Thoracic Diagnosis, Regeneration and Applied Immunology, Department of Thoracic Surgery, Medical University of Vienna; Aposcience AG, Vienna, Austria; Department of Plastic and Reconstructive Surgery, Medical University of Vienna, Vienna, Austria; Department of Medicine III, Division of Nephrology and Dialysis, Department of Internal Medicine III, Medical University of Vienna, Vienna, Austria; Department of Orthopedics and Trauma-Surgery, Medical University of Vienna, Vienna, Austria; Department of Dermatology, Medical University of Vienna, Vienna, Austria

**Keywords:** Scar, hypertropic scar, regeneration, wound healing, secretome, peripheral blood mononuclear cells

## Abstract

**Background:** Hypertrophic scars can cause pain, movement restrictions, and reduction of quality of life. Despite numerous options to tackle hypertrophic scarring, efficient therapies are still scarce, and cellular mechanisms are not well understood. Secreted factors from peripheral blood mononuclear cells (PBMCsec) were previously described for their beneficial effects in tissue regeneration. Here, we investigated the effects of PBMCsec on skin scarring in mouse models and human scar explant cultures at single cell resolution (scRNAseq).

**Methods:** Mouse wounds and scars were treated with PBMCsec either intradermally or topically. Human mature scars were treated with PBMCsec ex vivo in explant cultures. All experimental settings were analyzed by single cell RNA sequencing (scRNAseq). A variety of bioinformatics approaches were used to decipher gene regulation in the scRNAseq data sets. Components of the extracellular matrix (ECM) were investigated in situ by immunofluorescence. The effect of PBMCsec on myofibroblast differentiation and elastin expression was investigated by stimulating human primary fibroblasts with TGFβ.

**Findings:** Topical and intradermal application of PBMCsec regulated the expression of a variety of genes involved in pro-fibrotic processes and tissue remodeling. Our bioinformatics approach identified elastin as a common linchpin of antifibrotic action in both, the mouse and human experimental setting. *In vitro*, we found that PBMCsec prevents TGFβ-mediated myofibroblast-differentiation and attenuates abundant elastin expression through non-canonical signaling inhibition. Furthermore, TGFβ-induced breakdown of elastic fibers was strongly inhibited by addition of PBMCsec.

**Interpretation:** Together, we showed anti-fibrotic effect of PBMCsec on cutaneous scars in mouse and human experimental settings, suggesting PBMCsec as a novel therapeutic option to treat skin scarring.

**Research in context:** *Evidence before this study:* Paracrine factors secreted from irradiated peripheral mononuclear cells (PBMCsec) show strong tissue regenerative properties in a variety of organs and are shown to enhance cutaneous wound healing. Whether PBMCsec shows anti-fibrotic properties on scar formation has not been investigated so far.

*Added value of this study:* In the present study, we were able to demonstrate that PBMCsec improves quality of developing and mature scars in mouse and human scar tissue. We found that PBMCsec is able to attenuate the expression of various genes, promoting scar formation and inhibit TGFβ-induced myofibroblast differentiation. Elastin and TXNIP were identified as a common linchpin of its anti-fibrotic action.

*Implications of all the available evidence:* Using *in vivo, ex vivo*, and *in vitro* models and analyses on a single-cell level, our study paves the way for clinical studies evaluating the use of PBMCsec for the treatment of human cutaneous scars.

## Introduction

Skin scarring after surgery, trauma or burn injury is a major problem affecting 100 million people every year, causing a significant global disease burden (1). Patients with hypertrophic scars, occurring in 40-90% after injury (2), suffer from pain, pruritus, and reduced quality of life (3, 4). Skin scarring has been studied extensively (5, 6), and recently we were able to elucidate hypertrophic scar formation at a single-cell level (7). However, many cellular mechanisms still remain unclear, and most conservative therapeutic options have low evidence for their efficacy (8).Wound healing and scar formation are complex, rigidly coordinated processes, with multiple cell types involved (*9*). Wound healing is characterized by an acute inflammatory phase, a proliferative phase, and a remodeling phase (9). Prolonged inflammation results in increased fibroblast (FB) activity, with enhanced secretion of transforming growth factor beta 1 (TGFβ1), TGFβ2, insulin-like growth factor (IGF1), and other cytokines (10, 11). TGFβ1 induces the differentiation of FBs into myofibroblasts (myoFBs) (12). myoFBs show strong contractability and excessively deposit extracellular matrix (ECM)-components, eventually leading to (hypertrophic) scar formation. Matured (hypertrophic) scars show dense, parallel ECM and strong tissue contraction (12).

Numerous pharmaceutical attempts to tackle hypertrophic scars were proposed during the last decades, e.g. intralesional injection of corticosteroids, 5-Fluorouracil (5-FU), or triamcinolone (TAC) (13, 14). Other therapeutic options include compression therapy, or topical silicone application. These therapies, however, still lack evidence of efficacy and safety, show high recurrence rates, and mechanisms of action are not well understood (15, 16). In recent years, numerous pre-clinical studies suggested effective scar treatment or improvement of scar formation after application of conditioned media derived from different stem cell populations such as amnion mesenchymal stem cells (MSCs) (17), fat-derived stem cells (18), bone-marrow induced MSCs (19), induced pluripotent stem cells (20), amongst others (21). However, the transferability of promising pre-clinical animal studies to humans was shown to be limited (22). Furthermore, autologous conditioned media from various stem cell populations have significant disadvantages, as the production of these secretomes is expensive and hardly scalable, due to limited numbers of stem cells available(23).

Hence, the idea of cell-free paracrine therapies in an allogeneic setting has drawn increasing attention. As different kinds of stem cells still have the same limitations also in the allogeneic setting, peripheral blood mononuclear cells (PBMCs) were proposed as alternative source for paracrine factors (24).

The secretome of irradiated peripheral blood mononuclear cells (PBMCsec) was studied extensively in the past years and showed encouraging pre-clinical results. PBMCsec was found to enhance wound healing (25–27), elicit angiogenic effects (26, 28), prevent platelet aggregation and vasodilation (29), exert antimicrobial activity (30), attenuate neurological damage in focal ischemia (31) and spinal cord injury (32), and regenerate infarcted myocardium (33). Moreover, we found that PBMCsec was able to reduce activation of mast cells and basophils (34), reduce maturation and antigen uptake of dendritic cells as well as dendritic cell-mediated T-cell priming (35). Clinically, PBMCsec was found to be safe and well tolerated in topical application of autologous PBMCsec on skin wounds in a phase I study (36). In addition, a phase II clinical trial on the efficacy of allogeneic PBMCsec in patients with diabetic foot ulcers is currently ongoing (37). Of note, the favorable pleiotropic effects of PBMCsec cannot be broken down to a single mode of action (37), as PBMCsec was repeatedly demonstrated to exert its regenerative power by the synergistic action of all components, i.e. proteins, lipids, extracellular vesicles and nucleic acids (26, 28, 37).

Thus, we here attempt to provide a multi-model murine and human approach on a single-cell level to identify potential mechanisms of actions of PBMCsec on skin scarring. Due to the plethora of beneficial effects of PBMCsec, we hypothesize that PBMCsec prevents (hypertrophic) scarring or improves tissue quality in already persisting scars. Here we show anti-fibrotic activity of PBMCsec and provide mechanistic insights into its anti-fibrotic effect. The study facilitates the investigation of PBMCsec for its future clinical use as treatment option in skin scarring.

## Methods

### Ethics statement

The use of healthy abdominal skin (Vote Nr. 217/2010) and scar tissue (Vote Nr. 1533/2017) was approved by the Vienna Medical School ethics committee. Animal experiments were approved by the Medical University of Vienna ethics committee and by the Austrian Federal Ministry of Education, Science and Research (Vote Nr. BMBWF-66.009/0075-V/3b/2018).

### Patient material

Resected scar tissue was gained from three patients who underwent elective scar resection surgery after informed consent. Scars were previously classified as hypertrophic, pathological scar according to POSAS (38) by a plastic surgeon. All scars were mature scars, i.e. at least two years old, were not operated on and were previously not treated with corticosteroids, 5-FU, irradiation or similar. All scar samples were obtained from male and female patients younger than 45 years, with no chronic diseases or chronic medication. Healthy skin was obtained from three healthy female donors between 25-45 years from surplus abdominal skin removed during elective abdominoplasty.

### Animals

In all mouse experiments, 8-12 weeks old female Balb/c mice (Medical University of Vienna Animal Breeding Facility, Himberg, Austria) were used. Mice were housed according to enhanced standard husbandry at a 12/12h light/dark cycle in a selected pathogen-free environment, with food and water ad libidum.

### Full thickness wound and scarring model in mice

For a full thickness skin wound and scarring model, mice were deeply anesthetized with ketamine 80-100 mg/kg xylazine 10-12.5 mg/kg i.p., and postoperative analgesia by s.c. injection of 0,1ml/10mg Buprenorphin and Piritramid 7,5 mg/ml in drinking water. A 9×9mm square area was marked on the back and excised with sharp scissors. The wounds were left to heal uncovered without any further intervention for 4 weeks, and the resulting scar tissue was observed and fotodocumented.

### Production of irradiated mononuclear cell secretome (PBMCsec)

Secretome of human PBMCs were produced in compliance with good manufacturing practice (GMP) by the Austrian Red Cross, Blood Transfusion Service for Upper Austria (Linz, Austria) as described before (39, 40) (Figure S1). PBMCs were obtained by Ficoll-Paque PLUS (GE Healthcare, Chicago, IL, USA)-assisted density gradient centrifugation, adjusted to a concentration of 25×10^6^ cells/mL (25U/ml, 1Unit= secretome of 1 Million cells) and exposed to 60 Gy Caesium 137 gamma-irradiation (IBL 437C, *Isotopen Diagnostik CIS GmbH*, Dreieich, Germany). Cells were cultured in phenol red-free CellGenix GMP DC medium (CellGenix GmbH, Freiburg, Germany) for 24 ± 2 hours. Cells and cellular debris were removed by centrifugation and supernatants were passed through a 0.2 μm filter. For viral clearance, methylene blue treatment was performed as described (41). Secretome was lyophilized, terminally sterilized by high-dose gamma-irradiation, and stored at −80°C. All experiments were performed using secretomes produced under GMP of the following batches: A000918399086, A000918399095, and A000918399098, A000918399101, A000918399102, and A000918399105. Immediately before performing experiments, lyophilizate was resuspended in 0,9% NaCl to the original concentration of 25U/ml.

### PBMCsec injection of mouse scars

Starting on day 29 after skin wounding, mice were injected with 100yl 0,9% NaCl, medium (CellGenix GMP DC Medium phenol red-free), or PBMCsec, prepared as described above, every second day for two weeks. Subsequently, half of the mice from each group (n=2) were sacrificed and brought to analysis, the other half (n=2) were left for another two weeks without further intervention, and then sacrificed.

### PBMCsec topical application on mouse scars

Starting on the day of skin wounding (d0), mouse scars were treated with PBMCsec, medium or NaCl 0,9%. Ultrasicc/Ultrabas ointment (1:2; Hecht-Pharma, Bremervörde, Germany) was used as carrier substance for all treatments. Four parts Ultrasicc/Ultrabas and 1 part water were mixed and used as control treatment. For 5U/ml (200μl dissolved lyophilizate) PBMCsec or 200μl/ml medium were mixed with ointment. Mice were treated with control or inhibitors by applying 100□μl ointment on each wound immediately after wounding.

After application, mice were put individually in empty cages without litter for 30 min and monitored closely to prevent immediate removal of the treatments and allow sufficient tissue resorption. Scabs were left intact to prevent wound infections. Mice were treated daily for the first 7d, and thrice a week for 7 weeks. After scar formation, 4□mm biopsies of the scar tissue were taken and cut in half. One half of each scar sample was used for histological analysis, and the other biopsy halves from each treatment group were pooled and analyzed together with scRNAseq as described below.

### Ex vivo skin and scar stimulation

From human skin and scar tissue, 6mm punch biopsies were taken, subcutaneous adipose tissue was removed, and biopsies placed in 12-well plates supplemented with 400yl DMEM (Gibco, Thermo Fisher, Waltham, USA, with 10% fetal bovine serum and 1% penicillin/streptomycin), and 100yl CellGenix medium or 100yl PBMCsec. In addition, 100yl medium or PBMCsec were injected into upper dermis in the middle of the biopsy. Biopsies were incubated for 24h and then harvested for scRNAseq analysis. Sample “Skin 1 medium” was lost due to technical difficulties during preparation.

### Skin and scar PBMCsec stimulation, cell isolation and droplet-based scRNAseq

Mouse scars and human stimulated skin and scar samples were digested in Miltenyi Whole Skin dissociation Kit (Miltenyi Biotec, Bergisch-Gladbach, Germany) for 2.5h according to the manufacturer’s protocol and processed on a GentleMACS OctoDissociator (Miltenyi). The cell suspension was processed through a 100μm and a 40μm filter, centrifuged for 10min at 1500rpm, washed twice and resuspended in 0.04% FBS in phosphate buffered saline (PBS). DAPI 1μl/1 million cells was added for 30sec, cells were again washed twice and were sorted for viability on a MoFlo Astrios high speed cell sorting device (Beckman-Coulter, Indianapolis, USA), and only distinctly DAPI-negative cells were used for further processing. Immediately after sorting, viable cells were loaded on a 10X-chromium instrument (Single cell gene expression 3‘v 2/3, 10X Genomics, Pleasanton, CA, USA) to generate a Gel-bead in emulsion (GEM). GEM-generation, library preparation, RNA-sequencing, demultiplexing and counting were done by the Biomedical Sequencing Core Facility of the Center for Molecular Medicine (CeMM, Vienna, Austria). Sequencing was performed 2×75 bp, paired end on an Illumina HiSeq 3000/4000 (Illumina, San Diego, CA, USA).

### Cell-gene matrix preparation and downstream analysis

Raw sequencing reads were demultiplexed and aligned to human (GrChH38) and mouse (mm10) reference genome using Cell Ranger mqfast and count pipelines (v4.0, 10X Genomics, Pleasanton, CA, USA) to generate cell-gene matrices. Cell-gene matrices were then loaded into “Seurat” (v4.0, Satija Lab, New York, USA) in an R environment (v4.1.2, R foundation for statistical Computing, Vienna, Austria) and processed according to the recommended standard workflow for integration of several datasets (42, 43). All human skin and scar samples were integrated in a single integration, likewise, all mouse samples were integrated in a single integration. In samples, cells with less than 500 or more than 4000 detected genes, more than 20 0000 reads per cell or a mitochondrial gene count higher than 5% were removed from the dataset to ensure high data quality. After principal component analysis and identification of significant principal components by Jackstraw procedure (44), cells were clustered using non-linear dimensional reduction with Uniform Manifold Approximation and Projection (UMAP). Differentially expressed genes were calculated in Seurat using Wilcoxon rank-sum test with Bonferroni correction.

In all datasets, normalized count numbers were used for differential gene expression analysis, for visualization in violin plots, feature plots, dotplots, as recommended by guidelines(45). In all datasets, cell types were identified by well-established marker gene expression (Figures 2 and S5). For identification of differentially expressed genes (DEGs), normalized count numbers were used, including genes present in the integrated dataset to avoid calculation of batch effects. As keratin and collagen genes were previously found to contaminate skin biopsy datasets and potentially provide a false-positive signal (46), these genes (*COL1A1, COL1A2, COL3A1* and *KRT1 KRT5, KRT10, KRT14, KRTDAP*) were excluded from DEG calculation in non-fibroblast clusters (collagens) or non-keratinocyte clusters (keratins), respectively. Moreover, genes *Gm42418, Gm17056*, and *Gm26917* caused technical background noise and batch effect in mouse scRNAseq, as described before (47), and were thus excluded from the dataset.

### Gene ontology (GO)-calculation and dotplots

Gene lists of significantly regulated genes (adjusted *p*-value <0.05, average log fold change [avg_logFC] >0.1) were inputted in “GO_Biological_Process_2018” in the EnrichR package in R (v3.0, MayanLab, Icahn School of Medicine at Mount Sinai, New York, NY, USA). Dotplots were generated with ggplot2 (H. Wickham. ggplot2: Elegant Graphics for Data Analysis. Springer-Verlag New York, 2016.) with color indicating Adjusted.P.value and size showing the Odds.Ratio, sorted by adjusted p-value.

### GSEA-matrisome dotplots

Curated matrisome gene lists for the terms ‘NABA_ECM_GLYCOPROTEINS’, ‘NABA_COLLAGENS’, ‘NABA_PROTEOGLYCANS’, ‘NABA_ECM_REGULATORS’, and ‘REACTOME_ELASTIC_FIBRE_FORMATION’ were retrieved from the Gene Set Enrichment Analysis platform https://www.gsea-msigdb.org/gsea/index.jsp (48), and gene names were used to generate dotplots.

### TGFβ-injection fibrosis model in mouse skin

Mice were anesthetized with 3% isoflurane for three minutes. Mice were shaved in an approximately 1×1cm area intrascapularly. An approximately 4mm area was marked on the skin with permanent marker, and 800ng TGFβ1 dissolved in 100μl of NaCl 0,9%, Medium or PBMCsec (2,5U). Mice were injected with TGFβ1 and treatments in the marked area for 5 consecutive days and sacrificed on the sixth day. The marked injection areas were biopsied and prepared for histological analysis.

### Isolation of primary skin FBs

Primary skin and scar FBs were isolated as described previously (7). In brief, skin or scar samples were incubated in Dipase II (Roche, Basel, Switzerland) overnight. Subsequently, the epidermis was removed, and dermis was incubated in Liberase (Merck Millipore, Burlington, MA, USA) for two hours at 37°C. Afterwards, the tissue was filtered and rinsed with PBS, cells were plated in a T175 cell culture flask and cultured upon 90% confluency.

### Western Blots

Western blotting was performed as described previously (7). In brief, after lysis of cells in 1x Laemmli buffer, lysate was separated on SDS-PAGE gels (Bio-Rad Laboratories, Inc.), and proteins were transferred to nitrocellulose membranes and blocked with non-fat milk. After overnight incubation at 4° with primary antibody (Table of antibodies used, Figure S1B), membranes were incubated with horseradish-peroxidase conjugated secondary antibody and imaged.

### Immunofluorescence, H&E and EvG stainings

Immunofluorescence staining on formalin-fixed, paraffin-embedded (FFPE) sections of human and mouse skin and scar tissue were performed according to the protocol provided by the respective antibody manufacturer (Table of antibodies used, Figure S1B) as described previously (7). Hematoxylin and Eosin (H&E)-stainings and Elastica van Gieson (EvG)-stainings were performed at the Department of Pathology of the Medical University of Vienna according to standardized clinical staining protocols.

### TGFβ1-induced myofibroblast differentiation

TGFβ1-**s**timulation of primary FBs was performed as previously (7). Isolated primary FBs were plated in 6-well plates after the first passage and grown until 100% confluency. FBs were then stimulated with 10□ng/ml TGFβ1 (HEK-293-derived, Peprotech, Rocky Hill, NJ, USA), and with medium or PBMCsec for 24□h. Supernatants were removed and medium or PBMCsec were resupplied for another 24□h. Supernatants were collected and stored at −80□°C and cells were lysed in 1x Laemmli Buffer (Bio-Rad Laboratories, Inc., Hercules, CA, USA) for further analysis.

### Elastase assay

To measure elastase activity, a commercial kit (EnzChek® Elastase Assay Kit, E-12056, Thermo Fisher) was used according to the manufacturer’s instructions. Elastase was applied at 250mU/ml, and incubated with NaCl 0,9% (“Ctrl”), Medium or PBMCsec at 1:1 with assay buffer. Fluorescence intensity was measured with a BMG Fluostar Optima plate reader (BMG Labtech, Ortenberg, Germany) at 505/515nm wavelength (excitation/emission). Raw values were blank corrected and normalized to % of the averaged 4h of Ctrl samples. Samples were measured at 10min, 1h, 2h, 3h and 4h after elastase application. Statistical analysis was performed with a mixed-effects model for time factor, with Tukey’s multiple comparisons test.

### ELISA

Supernatants of TGFβ1-stimulated FBs after treatment with PBMCsec or controls were collected, centrifuged, and stored at −20□°C for further use. Protein levels of human Elastin ELISA (LS-F4567, LSBio, Seattle, USA) were measured according to the manufacturer’s manual. Absorbance was detected by FluoStar Optima microplate reader (BMG Labtech, Ortenberg, Germany).

## Results

### PBMCsec improves scar formation in mice after topical treatment during wound healing and intradermal injection of preformed scars

As our previous study on wound healing in pig burn wounds revealed a trend towards better tissue elasticity and less stiffness in early pig burn scars (27), we aimed to investigate the effect of PBMCsec on scar formation and on already existing scars in more detail on a single cell level.

Therefore, full-thickness excision wounds were set on the back of 6-8 weeks old female Balbc-mice and treated immediately by topical application of PBMCsec for 8 weeks (Figure 1A). In a second set of experiments, we let the scars develop for 4 weeks after wounding without further intervention and treated the formed scars by intradermal injection for 2 weeks. Scars were either analyzed directly after the two weeks of treatment or after another two weeks without further treatment to find out whether treatment-associated changes are permanent (Figure 1B).

**Figure 1:**
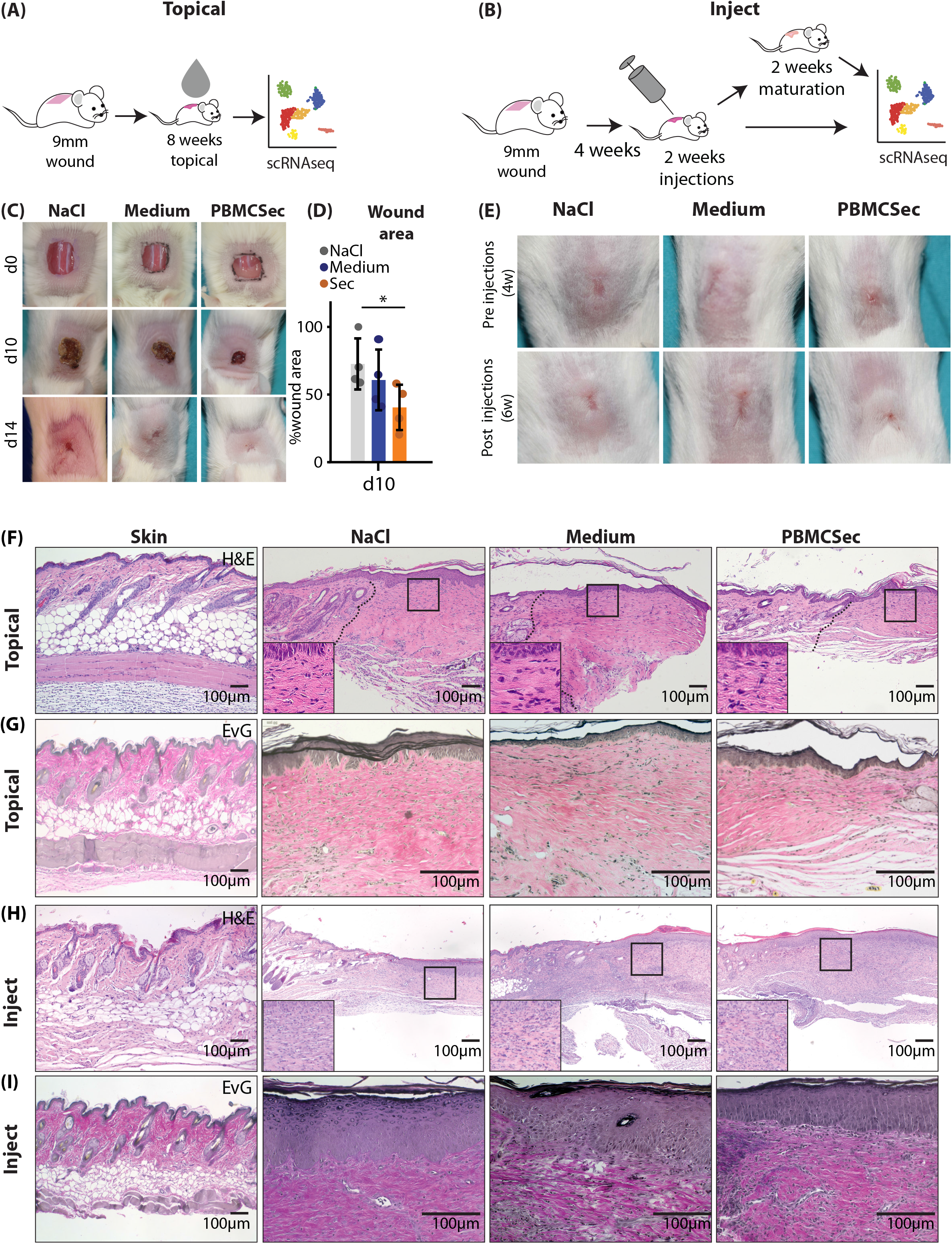
Comparison of PBMCsec-mediated effects in scars after topical or intradermal application. A, B) illustration “topical” or “inject” of workflow for mouse scars. A) 9×9mm square wounds were excised on mouse backs (n=4 per group), treated with topical PBMCsec for 8 weeks and brought to scRNAseq B) 9×9mm square wounds were excised on mouse backs (n=4 per group), left to mature for 4 weeks, injected with PBMCsec for 2 weeks and brought to scRNAseq, or matured for another 2 weeks and then brought to scRNAseq. C) Wound documentation of topically treated scars at post-wounding days 0, 10 and 14 D) Wound area measurements as normalized to d0 wound area of each respective wound E) Scar documentation of mouse scars before and after injections. F) Hematoxylin/eosin staining of ‘topical’ mouse wounds G) Elastica-van Gieson (EvG)-staining of ‘topical’ mouse wounds H) Hematoxylin/eosin staining of ‘inject’ mouse wounds I) Elastica-van Gieson (EvG)-staining of ‘inject’ mouse wounds.

As demonstrated previously with the secretome of non-irradiated PBMCs (25, 26) or PBMCsec in diabetic mice (25, 26), enhanced wound healing was also detectable in wild type mice after topical application of an emulsion containing PBMCsec (Figure 1C). PBMCsec reduced the wound size significantly stronger (40 +/-14% of the wound size) as compared to NaCl (72+/-16) and control medium alone (60+/-19%) (Figure 1D). Compared to intradermal injection of controls, scars appeared softer and reduced in size after injection of PBMCsec (Figure 1E). Histologically, scars showed a looser structure and reduced fiber density after topical PBMCsec as shown by hematoxylin/eosin (Figure 1F) and Elastica-van-Gieson (EvG) staining (Figure 1G). Of note, scars treated by intradermal injection showed a high number of infiltrating leukocytes (Figure 1H), presumably due to repeated tissue irritation from injections. However, the matrix was looser, and the orientation of collagen fibers showed more vertical structures after injection of PBMCsec (Figure 1I). Together these data indicate that PBMCsec not only improves wound healing but also scar formation and the quality of already existing scars in mice.

### PBMCsec induces significant changes of the transcriptome after topical and intradermal application

Next, we performed scRNAseq from scar tissue of the different experimental settings. After quality control (Figure S2A-C and E-G) clusters were defined according to well established marker genes (7, 49) from scRNAseq of topically treated scars (Figure S2D and H). Clusters aligned homogenously across all conditions (Figure 2A, E). Clusters were grouped in fibroblasts (FBs), smooth muscle cells and pericytes (SMC/PC), endothelial and lymphatic endothelial cells (EC), macrophages (Macro), Langerhans cells and dendritic cells (DC), T-cells and B-cells (TC), keratinocytes (KC), hair follicular cells (HF), melanocytes (Mel) and adipocytes (Adipo) (Figure 2A). Notably, one fibroblast cluster, FB 4, was expanded after topical treatment with PBMCsec compared to controls (Figure 2A,B), suggesting an important role in the anti-fibrotic action of PBMCsec.PBMCsec. Furthermore, relative numbers of DCs and TCs were increased with control medium, but slightly reduced with PBMCsec (Figure 2B). We then calculated differentially expressed genes (DEGs) of all cell populations of PBMCsec-treated scars compared to medium and NaCl-treated scars. Interestingly, significantly more genes were downregulated than upregulated after topical application of PBMCsec (Figure 2C), and the highest numbers of regulated genes were found in FB (red bars), macrophages (pale green bars) and KC (yellow bars) (Figure 2C) (35). To provide an impression of the overall regulation in all celltypes, the top 50 DEGs per clustergroup are shown in Figure S3A-I. Upregulation of numerous genes, previously described to be increased in scar tissue (7, 50, 51), was significantly inhibited after PBMCsec-application.

**Figure 2:**
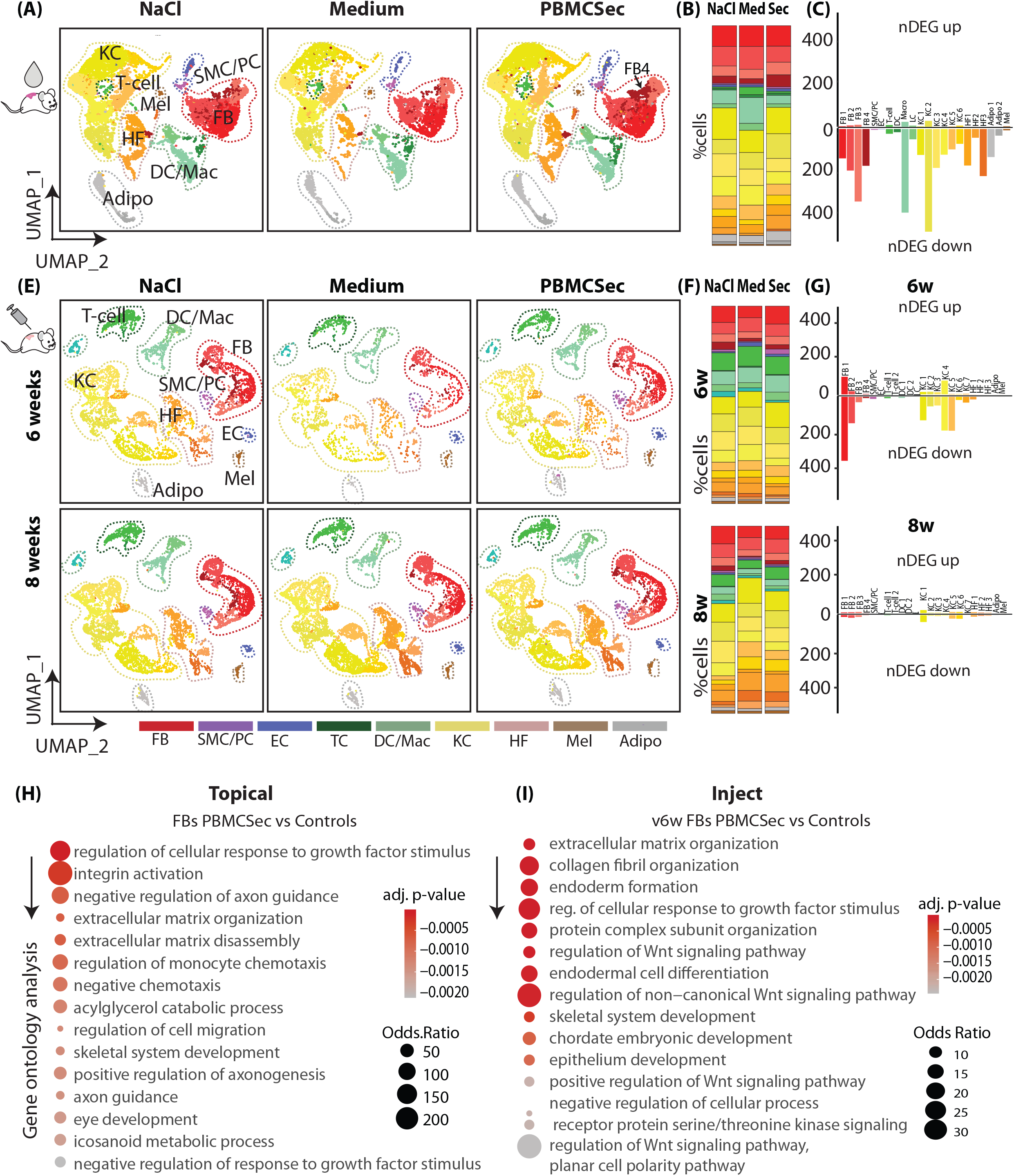
PBMCsec induces significant changes of the transcriptome after topical and intradermal application. A) UMAP-clustering of ‘topical’ mouse wounds (n=4 per condition, pooled for scRNAseq-analysis), split by condition, four fibroblast clusters (red, FB1-4), smooth muscle cells (SMC) and pericytes (purple, PC), endothelial cells (blue, EC), T cells (dark green TC), macrophages (Mac) and dendritic cells (light green, DC) three keratinocyte clusters (yellow, KC1-6), hair follicles (beige, HF 1-3) and melanocytes (brown, Mel), Adipocytes (grey). Clusters were grouped to ‘FB’, ‘PC’, ‘TC’, ‘DC’, ‘KC’, ‘HF’, ‘MEL’ and ‘Adipo’ for readability B) percentage of cells per cluster, split per condition C) number of significantly up- (y-axis positive) and downregulated (y-axis negative) genes (‘nDEG’) per cluster in ‘topical’ mice E) UMAP-clustering of ‘inject’ mouse wounds (n=2 per condition), split by condition. 6w = mice after two weeks injections, 8w =mice after injections + 2 weeks maturation; Clusters FB1-4, SMC, PC, EC, T cells1+2, DC1+2, KC1-7, HF 1-3, Mel), Adipo; Clusters were grouped to ‘FB’ (red), ‘PC’ (purple), ‘EC’ (blue), ‘TC’ (darkgreen), ‘DC’ (lightgreen), ‘KC’ (yellow), ‘HF’ (beige), ‘MEL’ (brown) and ‘Adipo’ (grey) for readability. F) percentage of cells per cluster, split per condition. G) Number of significantly up- (y-axis positive) and downregulated (y-axis negative) genes (‘nDEG’) per cluster in ‘inject’ mice, split by 6w and 8w H) Gene ontology (GO)-term calculation in ‘topical’ FBs of downregulated genes of PBMCsec compared Medium I) GO-term calculation in 6w ‘inject’ FBs of downregulated genes of PBMCsec vs Medium. DEGs were calculated per cluster comparing 8- vs 6-week-old scars using a two-sided Wilcoxon-signed rank test, including genes with average logarithmic fold change (avg_logFC) of >0.1 or <-0.1., adj. *p*-value <0.05. UMAP, uniform manifold approximation and projection.

Next, we similarly analyzed the scRNAsec dataset of scars which were treated by intradermal injection of PBMCsec and controls. After cluster identification and quality control (Figure S2D-G), clusters aligned homogenously across samples and conditions (Figure 2E). Whereas the cellular composition of the scars did not change after 6 weeks, the FB and immune cell populations were significantly reduced in 8 weeks old scars. (Figure 2F). Strikingly, there were again far more down-than upregulated genes also in the injected scars, and transcriptome changes were highest in FB1 and KC clusters (Figure 2G) after injections. Interestingly, only minor transcriptome changes remained in PBMCsec treated scars 8 weeks after wounding (Figure 2C). The top 50 DEGs after 6 weeks are shown per clustergroup in figure S4A-I. Remarkably, numerous genes regulated in the topical dataset and previously found relevant in skin scarring and mouse scar formation (7) were also regulated after PBMCsec injection (Figure S4A-I).

As the highest number of gene regulation was observed in FBs, and FBs are the main cell type involved in fibrotic processes, we further performed gene-ontology analysis of genes downregulated by PBMCsec-application in FBs in both experimental settings (Figure 2H and I). For topical application, our analysis revealed that genes downregulated by PBMCsec mainly show strong association with response to growth factors, integrin activation, monocyte chemotaxis and extracellular matrix organization, suggesting that activation of these processes is, at least partially, reduced (Figure 2H). GO term calculation from downregulated genes in FB after injection of PBMCsec revealed changes in ECM- and collagen organization, the response to growth factor stimulus and Wnt-signaling (Figure 2I).

Taken together, these bioinformatics data suggest an anti-fibrotic, anti-inflammatory effect of PBMCsec during scar formation, predominantly abrogating excessive matrix deposition.

### PBMCsec significantly alters the matrisome

Since FBs contributed most to transcriptome alterations by PBMCsec and GO-analysis indicated that genes associated with ECM were highly affected, we further assessed genes of the matrisome in more detail. Differentially regulated genes in all FBs after topical (Figure 3A-D) and intra-dermal injection (Figure 3E-H) were analyzed, using the curated matrisome gene set enrichment analysis (GSEA) gene lists (52). For better visualization, the whole matrisome was split in the main components collagens, proteoglycans, glycoproteins, and ECM regulators. Interestingly, most of the matrisome related genes were strongly downregulated by PBMCsec after topical and intradermal application (Figure 3). Similarly, most of the proteoglycans, glycoproteins and ECM regulators showed reduced expression after PBMCsec treatment. However, some of glycoproteins and ECM regulators, including *Fn1, Igfbp4/5, Ecm1, Postn*, and *Mfap5*, were even enhanced after PBMCsec treatment (Figure 3C-D), suggesting a targeted regulation of these factors.

**Figure 3:**
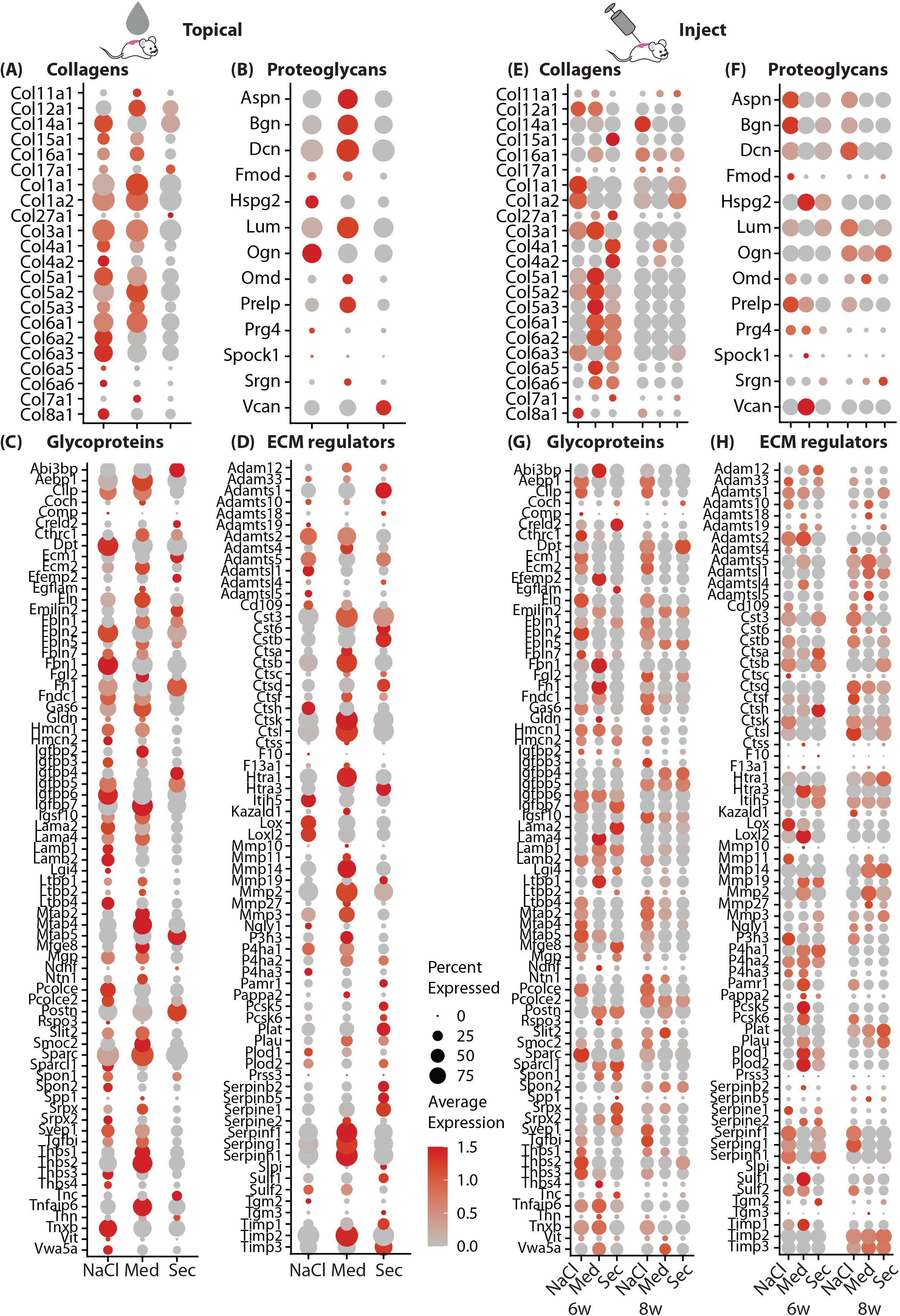
PBMCsec significantly alters the matrisome. Dotplots of gene lists of gene set enrichment of matrisome terms A, E) ‘collagens’, B,F) ‘proteoglykans’, C, G) ‘Glycoproteins’, D,H) ‘ECM-regulators inputted to FBs of the ‘topical’ (A-D) and ‘inject’ (E-H) datasets, split by condition. Circle size correlates with percent of cells expressing the respective gene, color (red) correlates with normalized fold change of expression.

Importantly, we also identified a variety of proteases, including *Mmp19* (matrix metalloprotease 19), *Ppcsk5/6* (Subtilisin/Kexin-Like Protease PC5/6) and *Adamts1* (A disintegrin-like and metallopeptidase with thrombospondin type 1 motif) to be regulated by PBMCsec.PBMCsec. Furthermore, the plasminogen activator/urokinase (*Plau*) and the plasminogen activator/tissue type (*Plat*) as well as the serine proteases *Htra1*, *Htra3* and *Aebp1* were elevated after topical application and intradermal injection of PBMCsec.PBMCsec. However, also a variety of protease inhibitors, including *Timp1 and 3* (Metallopeptidase Inhibitor 1 and 3), *Slpi* (Secretory Leukocyte Protease Inhibitor) and the potent urokinase inhibitors *Serpine1, Serpinb2* and *Serpinb5* were increased as well (Figure 3C,D). These findings confirm our previous work, highlighting the role of proteases and their inhibitors in skin fibrosis (7) and indicate that PBMCsec is able to interfere with the protease system contributing to scar formation.

### scRNAseq analysis of human skin and scars treated with PBMCsec ex vivo shows strong similarities to mouse models

As we showed an anti-fibrotic effect of PBMCsec during scar formation in mice, we next investigated its effect in human skin and ex vivo cultures of scar tissue. Therefore, we treated biopsies of human skin and human hypertrophic scars with medium or PBMCsec and cultivated them for 24h (Figure 4A). After quality control and cluster identification (Figure S5A-D), clusters aligned homogenously across donors and conditions (Figure 4B, Figure S5E). As described in our previous work, the ratio of FBs was increased in scars compared to skin (7), and several FB clusters, here clusters FB5 and FB7 were specifically found in scars (Figure 4B,C). Remarkably, the percentage of FBs, DCs and T-cells were reduced in scars after PBMCsec treatment (Figure 4C).

**Figure 4:**
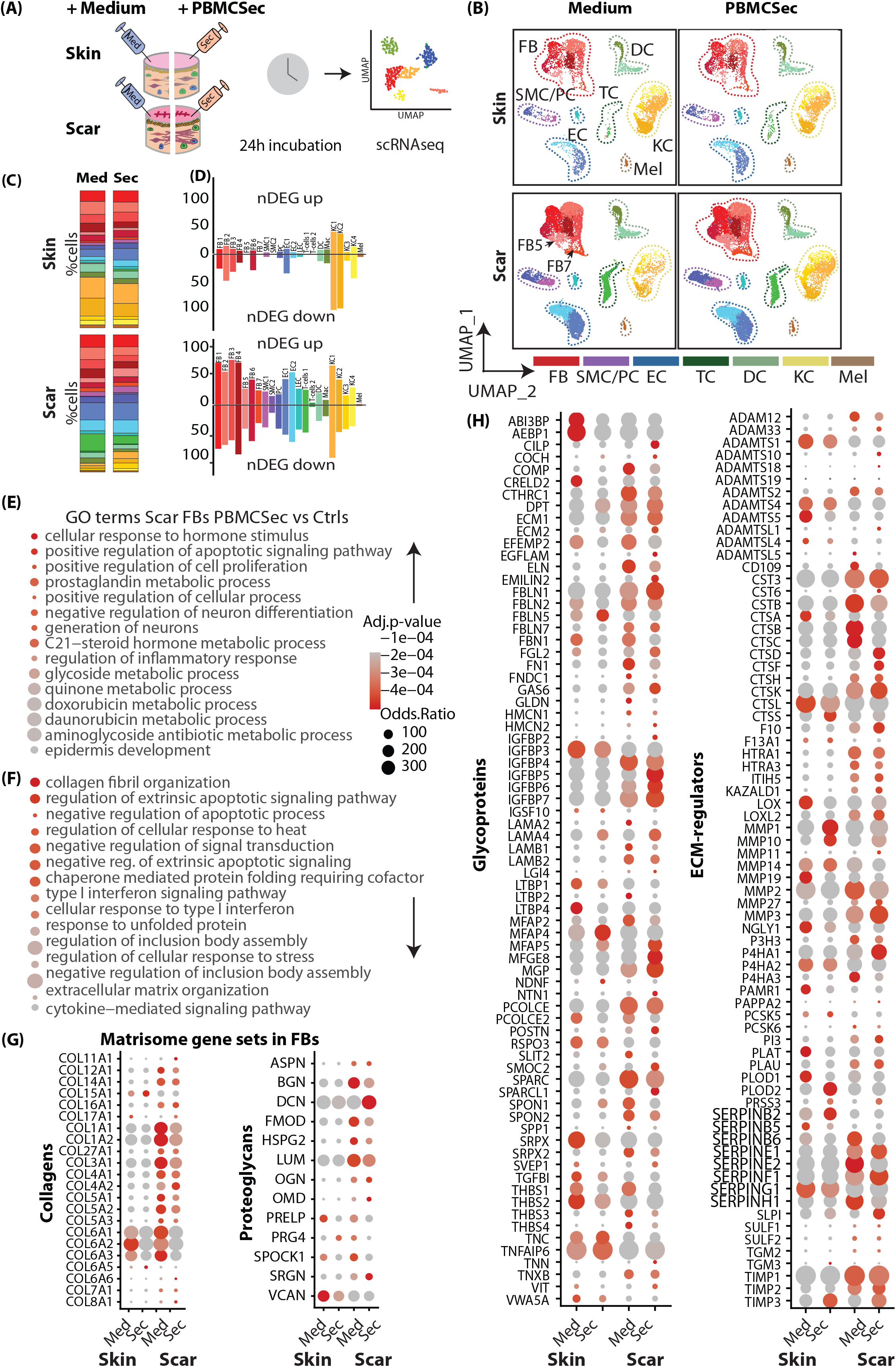
scRNAseq analysis of human skin and scars treated with PBMCsec ex vivo shows strong similarities to mouse models. A) illustration of scRNAseq-workflow in human skin and scar samples. Skin and scar biopsies were incubated overnight in medium or PBMCsec and brought to scRNAseq. B) UMAP-clustering of human skin and scar, split by condition. Seven fibroblast clusters (red, FB 1-7), smooth muscle cells (SMC) and pericytes (purple, PC), endothelial cells (blue, EC), T cells (dark green TC), macrophages (Mac) and dendritic cells (light green, DC) four keratinocyte clusters (yellow, KC1-4) and melanocytes (brown, Mel). Clusters were grouped to ‘FB’, ‘PC’, ‘TC’, ‘DC’, ‘KC’, ‘MEL’ and ‘HF’ for readability C) percentage of cells per cluster, split per condition D) number of significantly up- (y-axis positive) and downregulated (y-axis negative) genes (‘nDEG’) per E) Gene ontology (GO)-term calculation in ‘topical’ FBs calculated from E) upregulated and F) downregulated genes of PBMCsec compared to medium. G) Dotplots of gene lists of gene set enrichment of matrisome terms ‘collagens’, ‘proteoglykans’, H) ‘Glycoproteins’ ‘ECM-regulators’ inputted to FBs. DEGs were calculated per cluster comparing 8- vs 6-week-old scars using a two-sided Wilcoxon-signed rank test, including genes with average logarithmic fold change (avg_logFC) of >0.1 or <-0.1., adj. *p*-value <0.05. UMAP, uniform manifold approximation and projection.

Next, we calculated DEGs separately for skin (Figure S6) and scars (Figure S7) and found a much higher number of DEGs in scars as compared to normal skin, indicating a strong effect of PBMCsec on fibrotic tissue (Figure 4D). In line with our mouse datasets, most gene regulations were found in the FB clusters slightly more genes were downregulated than upregulated, particularly in skin (Figure 4D). Numerous genes we previously described for their regulation in hypertrophic scars (7) were also favorably regulated by PBMCsec (Figure S6, S7).

Next, we performed GO-term analysis from DEGs of FBs with PBMCsec compared to medium. In line with mouse data, downregulated terms (Figure 4F) included collagen fibril and ECM-organization, cytokine signaling pathway, and negative regulation of signal transduction, regulation of extrinsic apoptotic signaling pathway and type I interferon signaling pathway. Intriguingly, among upregulated terms (Figure 4E), negative regulation of neuron differentiation and generation of neurons was present. As we previously demonstrated that Schwann cells promote ECM-formation in keloids and affect M2 polarization of macrophages (53), this finding might hint a mechanism of PBMCsec affecting also this crosstalk.

Next, we assessed genes of the matrisome in the human data set (Figure 4H). Similarly, to the data obtained for mouse scars, collagens *COL1A1, COL3A1*, and *COL6A1/2/3* were also strongly downregulated, in scar more than in skin, and proteases *MMP1/MMP3/10* as well as protease inhibitors *SERPINE1/G1/F1/B2, SLPI* and *TIMP3* were upregulated (Figure 4D). Of note, PBMCsec increased the expression of *PI3* also in human scar tissue, an elastase-specific protease inhibitor, indicating a regulating effect not only in collagens, but also in elastic ECM components.

Together, our analysis in human *ex vivo* skin and scars corroborated findings of the *in vivo* mouse experiments, indicating an ECM-balancing, antifibrotic effect.

### PBMCsec abolishes myofibroblast-differentiation *in vitro*

After comprehensive analysis of the effects of PBMCsec in mouse and human models on single cell level, we next investigated the underlying mechanisms of the observed antifibrotic activity *in vitro*. Using a well-established *in vitro* fibrosis model (7, 54), we stimulated primary human skin FBs with TGFβ1 and investigated the effect of PBMCsec on myofibroblasts (myoFBs) formation (55). Upon stimulation of FBs with TGFβ1, FBs showed robust differentiation to αSMA-expressing myoFBs in all control treatments (NaCl and medium) (Figure 5A). In contrast, addition of PBMCsec completely abolished myoFB-dfferentiation and αSMA-expression (Figure 5A, B). As our scRNAseq revealed that of all major ECM components, *Eln/ELN* was most consistently downregulated in the matrisome of both mouse and human, we further assessed the effect of PBMCsec on the expression of elastin in vitro in FBs. Strikingly, elastin protein and mRNA expression were strongly downregulated by PBMCsec (Figure 5A, C) and secretion of ELN into the supernatant was significantly inhibited (Figure 5D). Next, we investigated whether PBMCsec contains TGFβ-inhibitors. Therefore, we used a HEK-cell based reporter assay to assess activity of canonical TGFβ1-signaling. While PBMCsec showed little if any TGFβ1 activity, addition of PBMCsec to active TGFβ1 does not inhibit canonical TGFβ1 activity (Figure S8A). These data indicate that PBMCsec does not inhibit MyoFB-differentiation by inhibiting Smad2/3-mediated TGFβ1-ativity. suggesting more downstream inhibitory or non-canonical action.

**Figure 5:**
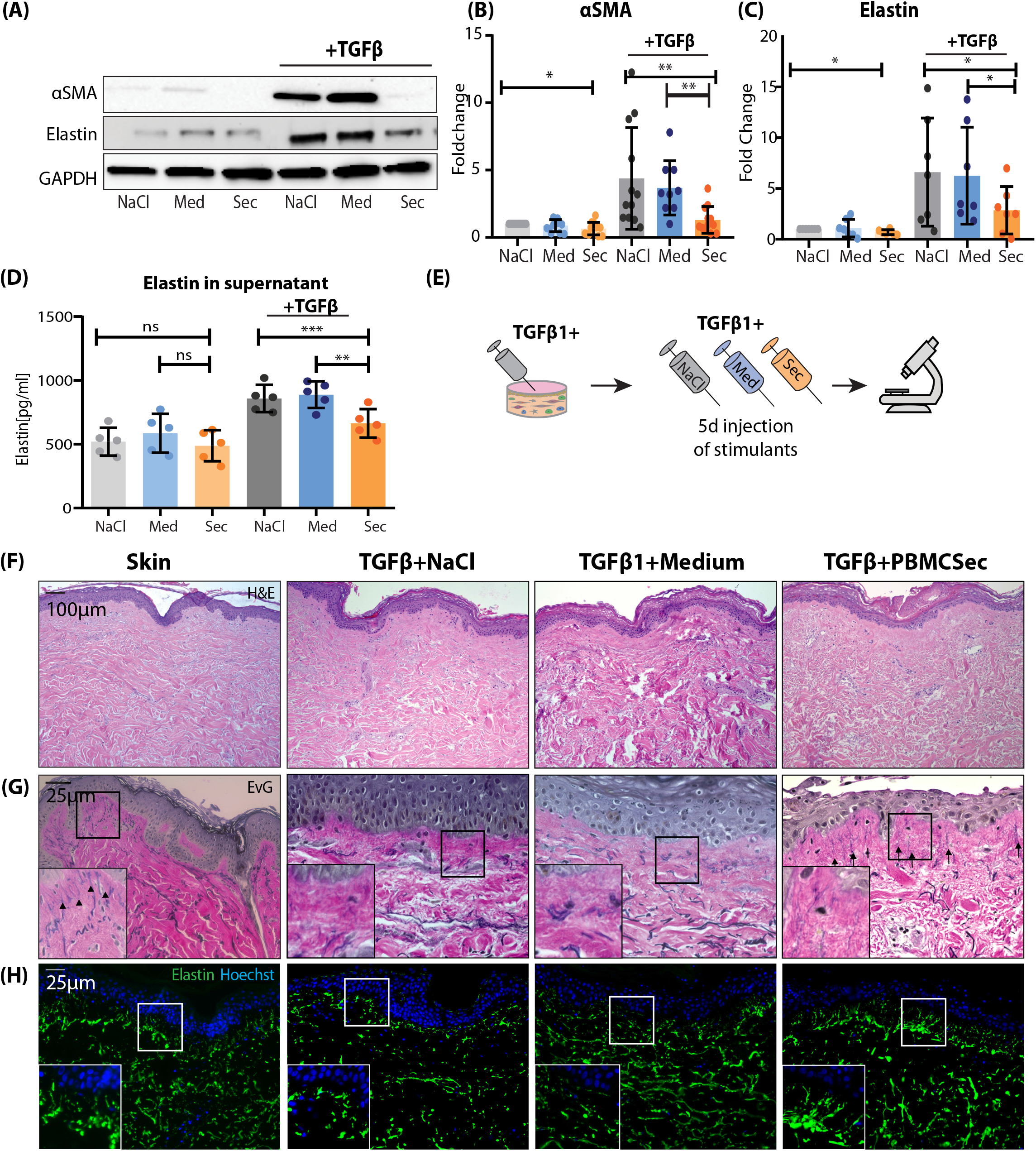
PBMCsec abolishes myofibroblast-differentiation *in vitro*. A) Western Blot stained for alpha Smooth muscle actin (SMA) and elastin, lysate from human primary FBs, stimulated with NaCl, Medium or PBMCsec, without or with TGFβ1, respectively B) Quantification of Western blot, normalized to ctrl (n=6 human donors) C) Elastin measured by ELISA from primary human FB supernatant, stimulated with NaCl, Medium or PBMCsec, without or with TGFß1 D) Workflow illustration of ex vivo human skin TGFß-stimulation experiment; 5mm skin biopsies were injected with TGFß and NaCl, Medium or PBMCsec for 5 consecutive days E) H&E F) Elastica van Gieson G) Immunofluoresence staining for elastin in human ex vivo skin samples. Statistical significance was tested using one-sided ANOVA. Lines and error bars indicate mean and standard deviation. NS *p*□>□0.05, \**p*□ <□0.05, \*\**p*□ <□0.01, \*\*\**p*□ <□0.001.

To corroborate the observed TGFβ-effects *in vivo*, we injected TGFβ1 into murine skin (modified after Thielitz et al. (54)) on 5 consecutive days (Figure S8B). While no morphological changes were observable in Hematoxylin-Eosin staining (Figure S8C), in immunostaining of Collagen I and III we found patches of increased matrix deposition in all samples (arrows, Figure S8D,E), that were not present in mice injected also with PBMCsec. Remarkably, we also observed accumulations of αSMA-expressing cells in the TGFβ1-injected deep murine dermis (squares), but not in PBMCsec-treated mice (Figure S8F).

Next, we aimed to further investigate changes in ECM-composition, especially elastin, in a human model. Thus, we injected TGFβ intra-dermally in human skin explants with and without NaCl, medium or PBMCsec (Figure 5E). Morphologically, no changes were observed in the H&E stainings (Figure 5F); however, when we stained for overall ECM-configuration using Elastica-van-Giesson staining (Figure 5G) and with immunofluorescence for Elastin (Figure 5H), we noticed specific subepidermal alterations in elastic fibers. In untreated skin, elastin showed vertical fibers reaching into the dermal papillae with parallel, horizontal fibers in the deeper dermis. These vertical, papillary fibers disappeared after TGFβ1-treatment, but were preserved when PBMCsec was added (Figure 5G, H). These data suggest that PBMCsec is able to reduce breakdown of elastic fibers, which occurs under after TGFβ stimulation.

### Combined analysis of murine and human scRNAseq datasets reveals Elastin and TXNIP as joint key players of beneficial PBMCsec-effects

To better understand the mutual mechanisms of action of ECM-balancing and anti-fibrotic mechanisms of PBMCsec, we performed a subclustering of FBs of all scRNAseq data sets (Figure S9D) and performed a combined analysis (Figure 6A). As myoFB, i.e. *Acta2/ACTA2-positive* FBs, disappear in mature scars (55), these cells were not detected in most of our data sets. Therefore, we were not able to investigate the effects of PBMCsec on myoFB-differentiation in our scar models in detail (Figure S9A-C). However, we detected a significant reduction of *ACTA2* in ex vivo PBMCsec-treated human scars (Figure S9C), indicating that even in mature scars, PBMCsec can reduce myoFB-content. When overlaying DEGs from FBs from all three experiments, no genes were mutually upregulated (Figure 6B). Interestingly, *Eln/ELN* and *Txnip/TXNIP* were mutually downregulated in all experimental settings (Figure 6C). Elastin and TXNIP were solidly reduced in all three scRNAseq, in both timepoint after injection, and in human scars (Figure 6D, E).

**Figure 6:**
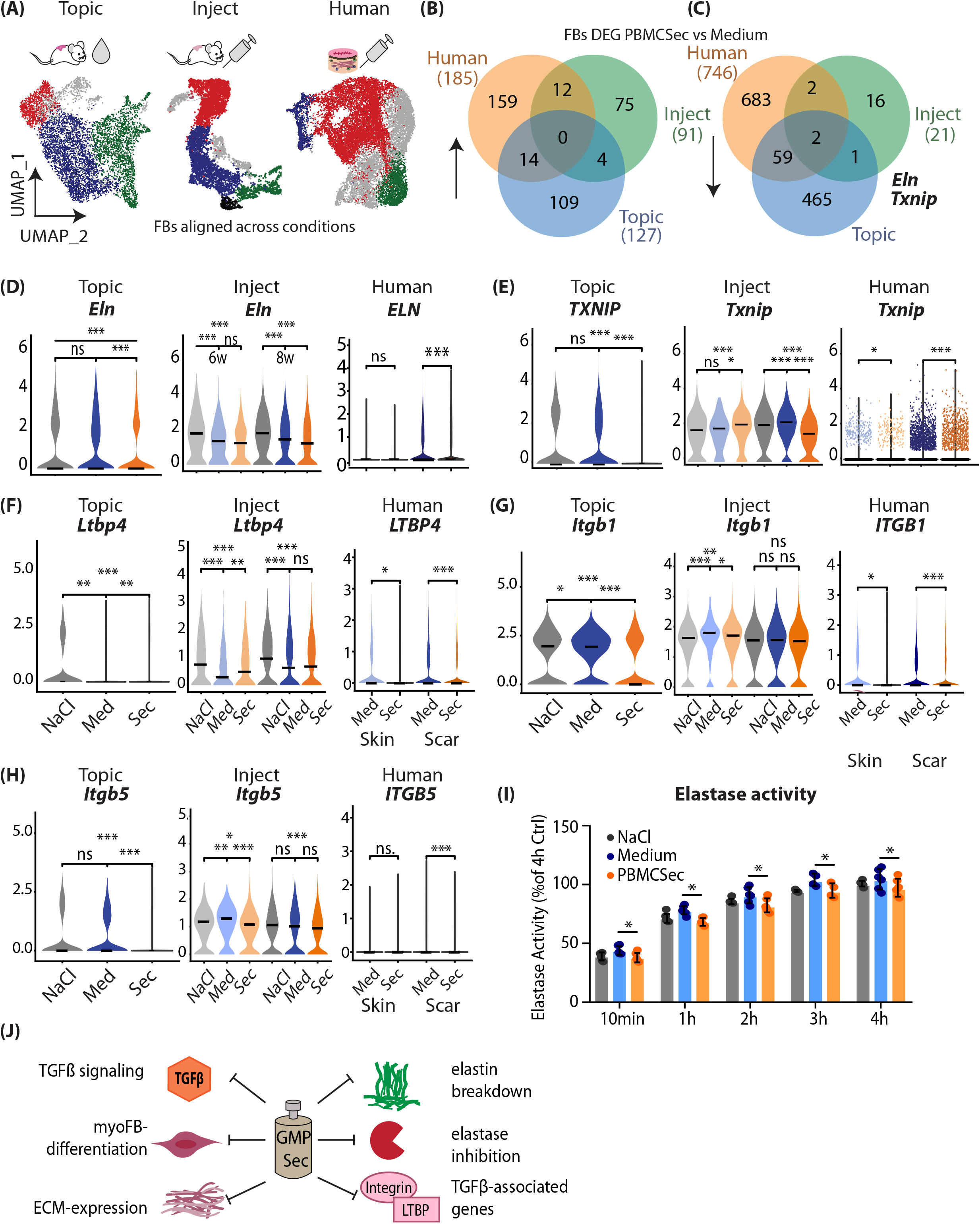
Combined analysis of murine and human scRNAseq datasets reveals Elastin as joint key player of beneficial PBMCsec-effects. A) Subclustering of FBs in ‘mouse topical’, ‘mouse inject’ and ‘human’ scRNAseq datasets and FB subcluster alignment. Red, cluster A; blue, cluster B; green, cluster C; aligned by cluster markers (Figure S9) B) Venndiagram of overlap of significantly upregulated and C) downregulated genes in FBs of all three datasets D-G) Violin plots of F) latent TGFβ binding protein 4 (Ltbp4/LTBP4) G,H) Integrin subunit beta 1/5 (*Itgb1/5/ITGB1/5*) in datasets. H) Elastase assay from fluorescence-marked pig pancreas elastase, with NaCl, Medium or PBMCsec added. Y-axis indicates fluorescence intensity, i.e. elastase activity. Comparison between groups was performed with Student t-test I) illustration of putative mechanisms of PBMCsec in scarrings In violin plots, dots represent individual cells, *y*-axis represents log2 fold change of the normalized genes and log-transformed single-cell expression. Vertical lines in violin plots represent maximum expression, shape of each violin represents all results, and width of each violin represents frequency of cells at the respective expression level. DEGs were calculated in FBs comparing Medium to FBs using a two-sided Wilcoxon-signed rank test, including genes with average logarithmic fold change (avg_logFC) of >0.1 or <-0.1 and Bonferroni-adjusted *p*-value <0.05. For violin plots, a two-sided Wilcoxon-signed rank test was used in R, NS *p*□ >□0.05, \**p*□ <□0.05, \*\**p*□ <□0.01, \*\*\**p*□ <□0.001.

As we have shown that PBMCsec does not interfere with canonical TGFB1 activity, we next wanted to know how TGFβ signaling is inhibited by PBMCsec.PBMCsec. TGFβ is one of the most pleiotropic signaling molecules, and interaction via regulation of its release and activation by elastin was described before (56). TGFβ is secreted inactive and bound to latent TGFβ binding proteins (LTBP1-4), together forming the large latent complex (LLC) (57). Activation of TGFβ occurs via a tightly controlled process, involving cleavage of the LTBPs, or protease-independent via integrins (57, 58). We therefore wondered, if PBMCsec also regulates molecules indirectly involved in TGFβ-activation. Surprisingly, we found that *Ltbp4/LTBP4* was decreased by PBMCsec in both mouse and human experimental settings (Figure 6F). *Ltbp4/LTBP4* is involved in both elastogenesis and regulation of TGFβ-signaling (58, 59), and increase of Ltbp4 is associated with fibrosis in scleroderma by TGF-β/SMAD signaling (60). Additionally, we found that expression of integrin subunits beta 1 and beta 5 (*Itgb/ITGB 1/5*) were also decreased upon PBMCsec treatment (Figure G,H). As both participate in the activation of TGFβ (57), these data indicate that their downregulation might indirectly contribute to the reduction of TGFβ-mediated fibrotic effects.

Finally, we investigated whether PBMCsec contains endogenous elastase inhibitors, which inhibit elastin breakdown and release of TGFβ (61), further enhancing the anti-TGFβ-feedback-loop by PBMCsec. However, elastase activity assay showed only a weak reduction of elastase activity after addition of PBMCsec (Figure 6H). We thus suppose a multi-effect model for attenuation of fibrosis by PBMCsec (Figure 6J): We observed a direct inhibition of TGFβ1 -mediated myoFB differentiation, but not via canonical signaling. Expression of numerous matrix genes is attenuated, and elastin secretion is significantly reduced. PBMCsec prevented elastin breakdown has mild elastase inhibition and interferes with TGFβ-induced gene expression (Figure 6J).

## Discussion

For a patient, a scar, particularly a hypertrophic scar, not only represents an aesthetic problem, but often also a significantly reduced quality of life, due to associated limitation of movement, itching and pain (8). As the treatment of hypertrophic scars is still difficult, the development of new therapeutic options is of particular interest. Here we presented a multi-model approach to assess the effects of a secretome-based drug (PBMCsec) on scar formation and scar treatment in mice and humans. Strong tissue-regenerative activity of PBMCsec has already been demonstrated not only in cutaneous wounds (25–27), but also in a variety of other organs, such as focal brain ischemia (31), spinal cord injury (32), and infarcted myocardium (33). Interestingly, in all organs mentioned before PBMCsec significantly reduced the size of the damaged areas and reduced the developing fibrotic tissue, suggesting a potential use for the treatment of cutaneous scars (33)*, 27*). In this study, we compared the effect of PBMCsec on scar formation in mice *in vivo* and in human *ex vivo* explant cultures. In mice we performed intradermal injection of the secretome into mature scars and applied it topically during wound healing and scar formation. So far, just a few studies investigated the effects of paracrine factors on cutaneous scarring by using cell secretomes from different stem cell types, including umbilical cord stem cells, adipose tissue-derived stem cells or mesenchymal stem cells (62–64). Arjunan et al. and Liu et al. showed that conditioned medium from umbilical cord Wharton’s jelly stem cells or adipose tissue-derived stem cells reduced the activation and growth of keloidal fibroblasts *in vitro* and in *in vivo* keloid models (63). In addition, Hu et al. suggested a combined treatment of conditioned medium from MSC and botulinum toxin for the treatment of hypertrophic scars (63). However, in depth analyses of the underlying mechanisms are still lacking. Hitherto, there are no studies available, investigating the effect of cell secretomes on scar formation using a holistic approach. Our study is the first to use scRNAseq to unravel mechanisms, important for improved scar formation after application of a cell secretome.

Interestingly, in our mouse experiments, both application routes, topical and intradermal application, showed promising effects on scar formation and scar treatment. Of note, significantly more genes were regulated after topical application of PBMCsec, suggesting higher efficacy of topical application in wounds than after injections. However, the improved wound healing process per se after PBMCsec application might already be decisive for a better scar quality. Therefore, a direct comparison of the two application routs is difficult, and needs further experiments, in which PBMCsec has to be applied topically on already existing scars. Furthermore, potential other treatment options, such as application after laser treatment (65), microneedling (66, 67), or together with nanocarriers (67) should be tested in future experiments. Most importantly, and in line with the data on mouse scar formation, we also identified a significant anti-fibrotic effect of PBMCsec on human mature hypertrophic scars in explant cultures. In fact, the treatment of scars with PBMCsec in mice and humans showed high similarities. In both species we found the strongest transcriptome alterations in FB clusters, specifically in genes of the so called matrisome, i.e. collagens, proteoglycans, glycoproteins, and ECM regulators (22–24). The matrisome, developed for large-scale *in silico* analyses, has been defined recently and provides a comprehensive overview on components of the ECM (52, 68, 69). Although several characteristics of ECM alterations in (hypertrophic) scars have already been described (70), our study provides the first big data set analyzing changes of the entire matrisome in mice and humans during wound healing and scar formation. These highly valuable data set will be the basis for many future studies on the pathophysiology of wound healing and scar formation, as well as on the effects of secretome-based scar treatment.

In the present study, we further focused on elastin, which was similarly downregulated by PBMCsec in all conditions and species investigated. Elastin fibril sequences interact with microfibrils and bind to cell surface receptors (71). Elastin is extremely durable and has a half-life of ~70 years (71, 72). While intact elastin is inert and insoluble, it can be degraded by a plethora of elastases (72), including MMPs, aspartic proteases, serine proteases and cysteine proteases (72). In our *ex vivo* assays we found strong degradation of elastic fibers in human skin by TGFβ, which was completely inhibited by PBMCsec, suggesting an elastase inhibiting effect of PBMCsec.PBMCsec. Intriguingly, this effect of TGFβ on elastic fibers appears counterintuitive and we did not find any other study describing this phenomenon. The interaction of TGFβ and elastin is complex. TGFβ is generally known to induce elastogenesis (48), stabilize elastin mRNA (48, 49) and increase elastin secretion (Figure 5), which is most likely due to post-transcriptional control of elastin (48). This is in line with our *in vitro* finding, where we could show strong upregulation of elastin production in fibroblasts treated with TGFβ. Interestingly, also this upregulation was significantly inhibited by PBMCsec on the mRNA and protein level. So far, we cannot offer an explanation for this phenomenon. It is tempting to speculate that the proteolytic breakdown of elastin triggers de novo synthesis of elastin.

Furthermore, whether TGFβ-induced over-production of elastin also leads to the assembly of new functional elastic fibers is still not fully understood. Therefore, the mechanisms by which PBMCsec inhibits elastin breakdown needs further investigations. Interestingly, our *in vitro* elastase assay showed only weak anti-elastase activity of PBMCsec, suggesting that either the specific enzyme inhibited by PBMCsec is not detected by the *in vitro* assay, or PBMCsec leads to an induction of endogenous protease inhibitors. In line with the second hypothesis, Copic et al. recently showed that PBMCsec is indeed able to induce the production of SERPINB2, a serine protease inhibitor, in human mononuclear cells [REF].(73). Furthermore, with scRNAseq, we showed that some elastase inhibitors, such as *PI3* (peptidase inhibitor 3) and *SLPI* (secretory leukocyte protease inhibitor) were significantly upregulated in FBs by PBMCsec in scars (Figure 4H). Despite being well investigated for their beneficial effects in cystic fibrosis (74), these elastase inhibitors have been hardly assessed for their role in cutaneous scar formation so far. Further, more sophisticated experiments are needed to fully address the role of these enzyme inhibitors in scar formation.

Aside from elastin, the only other gene consistently regulated by PBMCsec in all three scRNAseq experimental approaches was *TXNIP* (Thioredoxin interacting protein). TXNIP is critically involved in the regulation of reactive oxygen species (ROS) and cellular oxidative stress (75), and was shown to contribute to disturbed wound healing under ischemic conditions (76). With regard to scar formation, TXNIP was shown to be elevated in a murine model of pulmonary fibrosis, and inhibition of TXNIP in this model led to a reduction of ROS and myoFB differentiation (77). The exact role of TXNIP in skin pathologies and in scars, however, is scarcely investigated (78). Our finding that the downregulation of *TXNIP* is conserved across all our experimental approaches, suggests that PBMCsec-induced *TXNIP* downregulation might be an important mechanism, contributing to the anti-fibrotic action of PBMCsec.PBMCsec. However, further studies are needed to fully decipher the mechanism of TXNIP-regulation as well as its impact on cutaneous scar formation.

Interestingly, PBMCsec also prevented FB activation and myoFB differentiation. In line with our results, previous studies showed that treatment of FBs with conditioned medium of mesenchymal or pluripotent stem cells were able to reduce myoFB-differentiation (79, 80). In contrast to these studies, we were not able to identify a direct inhibitory action of PBMCsec on canonical TGFβ/Smad signaling (80). However, TGFβ has been shown to also induce fibrosis via non-canonical (non-SMAD) signaling pathways (81), and blocking non-canonical signaling prevents profibrotic phenotypes (82). Possible non-canonical pathways might include glycogen synthase kinase-3β (GSK-3β) (82), a pathway we previously found regulated upon non-SMAD TGFβ-mediated abolishment of myoFB-differentiation (7). Hitherto, only few secreted molecules inhibiting non-canonical TGF-signaling have been described. Del-1 (Developmentally-Regulated Endothelial Cell Locus 1 Protein) was shown to inhibit TGFβ and attenuate fibrosis via suppressing α_v_ integrin-mediated activation of TGFβ (83). In addition, several proteins, such as fibroblast growth factor (FGF), epidermal growth factor (IGF), interferon gamma and IL-10, all of which present in PBMCsec, are known to inhibit myoFB-differentiation (6). To identify the exact pathway of TGFβ-inhibition by PBMCsec, a detailed proteomic approach and assessment of multiple pathways will be necessary in the future.

Together, we provide an extensive study with multiple experimental approaches and ample scRNAseq data. Comprehensive analyses suggest a solid anti-fibrotic, ECM-reducing and myoFB-inhibiting effect of PBMCsec. We identified the prevention of elastin break-down as a putative major underlying mechanism of PBMCsec-mediated scar attenuation. We thus propose future clinical assessment of PBMCsec to attenuate skin scarring during wound healing, and to treat already existing mature scars.

## Contributors

VV, HJA, and MM: design of the study. VV, DC, KK, MD and BG: data acquisition. VV, DC, KK, MD, BG, HJA, and MM: data analyses and interpretations. CR, HJA and MM: resources. VV, DC, KK, MD and MM: access to the data and data verification VV, HJA and MM: drafting original manuscript. All authors: review and approval of the final manuscript.

## Data sharing statement

The scRNAseq data generated in this study have been deposited in the NCBI GEO database under accession numbers GSE156326 and GSE202544. The raw sequencing data are protected and are not available due to data privacy laws. If raw sequencing data are absolutely necessary for replication or extension of our research, they will be made available upon request to the corresponding author within a 2-week timeframe. All other relevant data supporting the key findings of this study are available within the article and its Supplementary Information files or from the corresponding author upon reasonable request.

## Acknowledgements

We thank Dr. HP Haselsteiner and Karl Fister, head of the CRISCAR Familienstiftung for their belief in this private public partnership to augment basic and translational clinical research. The authors acknowledge the core facilities of the Medical University of Vienna, a member of Vienna Life Science Instruments. This research project was finance in part by the Austrian Research Promotion Agency grant “APOSEC” (852748 and 862068, 2015-2019), by the Vienna Business Agency “APOSEC to clinic,” (2343727,2018-2020), and by the Aposcience AG under group leader HJA. MM was funded by the Sparkling Science Program of the Austrian Federal Ministry of Education, Science and Research (SPA06/055).

## Supplementary figure legends

**Figure S1: Methods**

A) Workflow of GMP-PBMCsec-production: leukocyte cones are obtained as blood-donation by-product, PBMCs are isolated by Ficoll-centrifugation, cells are irradiated with 60Gy and incubated for 24h. Superantatants are filtrated and lyophilized, and off-the-shelf PBMCsec is stored at −20° until use.

B) Table of antibodies used. IF = immunofluorescence, WB = western blot

**Figure S2: Quality control of mouse scRNAseq**

A-C, E-G) Violin plots of quality control parameters of the ‘topical’ and ‘inject’ dataset. A, E) total molecules per cell B, F) (gene count per cell and C, G) mitochondrial gene content. D, H) Feature Plots of cluster markers for cluster identification in the ‘topical’ and ‘inject’ dataset: *Col1a1* (collagen I alpha 1) for fibroblasts, *Acta2* (smooth muscle actin) for smooth muscle cells and myofibroblasts, *Rgs5* (Regulator Of G Protein Signaling 5) for pericytes, *Pecam* (Platelet And Endothelial Cell Adhesion Molecule 1) for endothelial cells, *Lyve1* (Lymphatic Vessel Endothelial Hyaluronan Receptor 1) for lymphatic endothelial cells, *Cd207* (Langerin) for Langerhans cells, *Cd3d* (cluster of differentiation 3D) for T-cells, *Itgax* for dendritic cells, *Aif1* (allograft inflammatory factor 1) for macrophages, *Mki67* (Marker Of Proliferation Ki-67) for proliferating cells, *Krt1* (Keratin1) for spinous and granular keratinocytes (KCs), *Krt5* (Keratin 5) for basal KCs, *Krt25* (Keratin 25) for hair follicles, *Pmel* (Premelanosome Protein) for melanocytes, *Cidea* (Cell Death Inducing DFFA Like Effector A) for adipocytes. Vertical lines in violin plots represent maximum expression, shape of each violin represents all results, and width of each violin represents frequency of cells at the respective expression level. In feature plots, normalized log expression of the respective gene is mapped onto the UMAP-plot. Color intensity indicates level of gene expressions. UMAP, uniform manifold approximation and projection.

**Figure S3**: **Top 50 regulated genes per cell group in ‘Topical’ mouse scars**

In A) fibroblasts (FBs, red circles), B) smooth muscle cells and pericytes (SMC/PCs, purple), C) endothelial cells (ECs, blue), D) T-cells (dark green), E) dendritic cells and macrophages (DCs, light green), and keratinocytes (KCs, yellow), hair follicles (HF, beige), melanocytes (Mel, brown), adipocytes (Adipo, grey); differentially expressed genes (DEGs) were calculated comparing ‘PBMCsec’ mouse scars to ‘NaCl’-mouse scars, using Wilcoxon rank sum test, including genes with average logarithmic fold change (avglogFC) of >0.1 or < −0.1 and Bonferroni-adjusted p-value <0.05. For each cellgroup, top 50 DEGs according to lowest adjusted p-value are displayed, split by treatment. Dot size represents percent of cells expressing the respective gene, color correlates with average expression.

**Figure S4**: **Top 50 regulated genes per cell group in 6 weeks ‘Inject’ mouse scars**

In A) fibroblasts (FBs, red circles), B) smooth muscle cells and pericytes (SMC/PCs, purple), C) endothelial cells (ECs, blue), D) T-cells (dark green), E) dendritic cells and macrophages (DCs, light green), and keratinocytes (KCs, yellow), hair follicles (HF, beige), melanocytes (Mel, brown), adipocytes (Adipo, grey); differentially expressed genes (DEGs) were calculated comparing ‘PBMCsec’ mouse scars to ‘NaCl’-mouse scars, using Wilcoxon rank sum test, including genes with average logarithmic fold change (avglogFC) of >0.1 or < −0.1 and Bonferroni-adjusted p-value <0.05. For each cellgroup, top 50 DEGs according to lowest adjusted p-value are displayed, split by treatment and timepoint. Dot size represents percent of cells expressing the respective gene, color correlates with average expression.

**Figure S5: Quality control of mouse scRNAseq**

A-C) Violin plots of quality control parameters of the ‘human’ dataset. A) total molecules per cell B) (gene count per cell and C) mitochondrial gene content. D) Feature Plots of cluster markers for cluster identification in the ‘topical’ and ‘inject’ dataset: *COL1A1* (collagen I alpha 1) for fibroblasts, *ACTA2* (smooth muscle actin) for smooth muscle cells and myofibroblasts, *RGS5* (Regulator Of G Protein Signaling 5) for pericytes, *SELE* (Selectin E) for endothelial cells, *LYVE1* (Lymphatic Vessel Endothelial Hyaluronan Receptor 1) for lymphatic endothelial cells, *Cd3d* (cluster of differentiation 3D) for T-cells, *ITGAX* for dendritic cells, *CD207* (Langerin) for Langerhans cells, *AIF1* (allograft inflammatory factor 1) for macrophages, *KRT1* (Keratin1) for spinous and granular keratinocytes (KCs), *KRT5* (Keratin 5) for basal KCs, *MLANA* (Melan-A) for melanocytes. E) UMAP-clustering in the ‘human’ dataset, split by samples. Sample “Skin 1 medium” was lost due to technical difficulties during preparation.

Vertical lines in violin plots represent maximum expression, shape of each violin represents all results, and width of each violin represents frequency of cells at the respective expression level. In feature plots, normalized log expression of the respective gene is mapped onto the UMAP-plot. Color intensity indicates level of gene expressions. UMAP, uniform manifold approximation and projection

**Figure S6: Top 50 regulated genes per cell group in human skin**

In A) fibroblasts (FBs, red circles), B) smooth muscle cells and pericytes (SMC/PCs, purple), C) endothelial cells (ECs, blue), D) T-cells (dark green), E) dendritic cells and macrophages (DCs, light green), and keratinocytes (KCs, yellow), melanocytes (Mel, brown); differentially expressed genes (DEGs) were calculated comparing ‘PBMCsec’ skin biopsies to ‘Medium’ skin biopsies, using Wilcoxon rank sum test, including genes with average logarithmic fold change (avglogFC) of > 0.1 or < −0.1 and Bonferroni-adjusted p-value <0.05. For each cellgroup, top 50 DEGs according to lowest adjusted p-value are displayed, split by treatment. Dot size represents percent of cells expressing the respective gene, color correlates with average expression.

**Figure S7: Top 50 regulated genes per cell group in human scar**

In A) fibroblasts (FBs, red circles), B) smooth muscle cells and pericytes (SMC/PCs, purple), C) endothelial cells (ECs, blue), D) T-cells (dark green), E) dendritic cells and macrophages (DCs, light green), and keratinocytes (KCs, yellow), melanocytes (Mel, brown); differentially expressed genes (DEGs) were calculated comparing ‘PBMCsec’ scar biopsies to ‘Medium’ scar biopsies, using Wilcoxon rank sum test, including genes with average logarithmic fold change (avglogFC) of > 0.1 or < −0.1 and Bonferroni-adjusted p-value <0.05. For each cellgroup, top 50 DEGs according to lowest adjusted p-value are displayed, split by treatment. Dot size represents percent of cells expressing the respective gene, color correlates with average expression.

**Figure S8: The interaction of TGFβ and PBMCsec *in vivo***

A) In vitro TGFβ-activity assay. HEK-cells colorimetrically detecting SMAD2/3 TGFβ-activity were incubated with recombinant TGFβ1, with PBMCsec alone, and with TGFβ1 and PBMCsec combined. Color intensity correlates with TGFβ 1-signaling activity. B) Workflow of mice intradermally injected with TGFβ1 and treatmens. Mice were intradermally injected with 800ng TGFβ1 dissolved 100μl in NaCl 0,9%, Medium or PBMCsec on five consecutive days and sacrificed on the 6^th^ day. C) H&E staining of resulting “scars” of the injected area. Immunofluorescence stainings of D) collagen 1, E) collagen 3, F) smooth muscle actin in TGFβ1-injected mouse skin. Arrows indicate areas of dense matrix deposition.

**Figure S9: Subcluster analysis of FB populations**

A-C) Violin plots of *Acta2/ACTA2* in mouse and human datasets, split by treatments. D) Feature plots of FB subcluster markers in FB subclusters of mouse and human datasets. *Tagln/TAGLN*, Transgelin; *Dpp4/DPP4*, dipeptidyl-peptidase 4; *Sfrp2/SFRP2*, Secreted Frizzled Related Protein 2; *Mgp/MGP*, Matrix Gla Protein; *Plau/PLAU*, urokinase; *Tgfbi/TGFBI*, Transforming Growth Factor Beta Induced. In violin plots, dots represent individual cells, *y*-axis represents log2 fold change of the normalized genes and log-transformed single-cell expression. Vertical lines in violin plots represent maximum expression, shape of each violin represents all results, and width of each violin represents frequency of cells at the respective expression level. In feature plots, normalized log expression of the respective gene is mapped onto the UMAP-plot. Color intensity indicates level of gene expressions. UMAP, uniform manifold approximation and projection. A two-sided Wilcoxon-signed rank test was used in R. NS *p*□>□0.05, \**p*□<□0.05, \*\**p*□ <□0.01, \*\*\**p*□ <□0.001.

