## Supplementary material for "The secretome of irradiated peripheral mononuclear cells attenuates hypertrophic skin scarring"

Figure S1

(A)

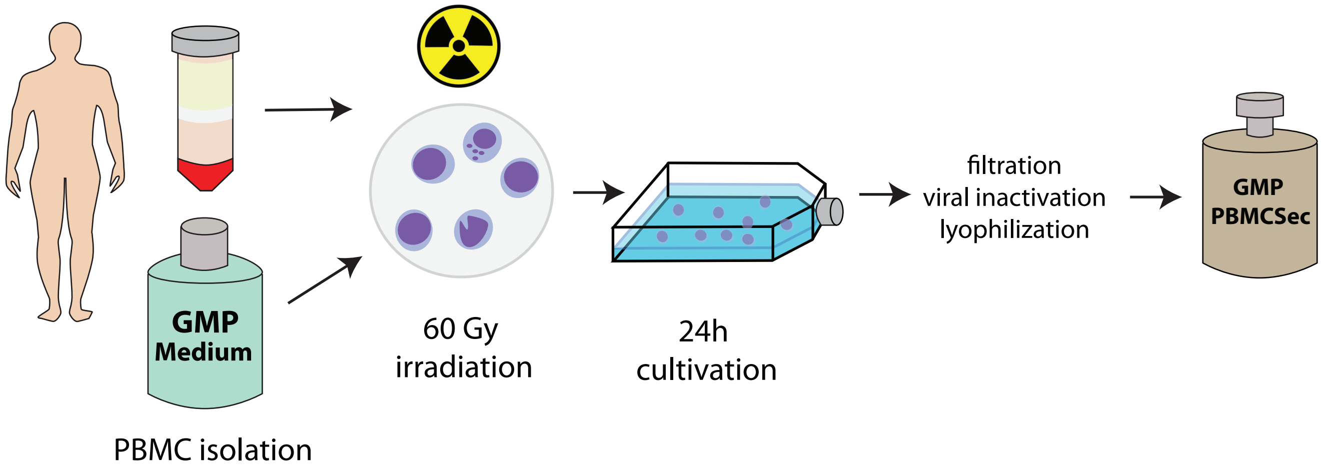


(B)

| Target | Supplier | Product Nr. | Host species | Dilution | Application |
|---|---|---|---|---|---|
| GAPDH | abcam | ab8245 | mouse monclonal | 1:10000 | WB |
| Collagen I | abcam | ab34710 | rabbit polyclonal | 1:200 | IF |
| Collagen III | abcam | ab7778 | rabbit polyclonal | 1:200 | IF |
| Elastin | Merck | Mab2503 | mouse monclonal | 1:100 | IF, WB |
| Alexa fluor® 546 anti-mouse IgG (H + L) | Invitrogen | A-11030 | goat polyclonal | 1:500 | IF, 2nd step |
| Alexa fluor® 546 anti-rabbit IgG (H + L) | Invitrogen | A-11035 | goat polyclonal | 1:500 | IF, 2nd step |
| Anti-mouse, HRP-conjugated | GE Healthcar | GENX-A931 | goat polyclonal | 1:10 000 | WB, 2nd step |
| Anti-rabbit, HRP-conjugated | Bio-Rad | #1706511 | goat polyclonal | 1:10 000 | WB, 2nd step |

**Figure S1: Methods**

A) Workflow of GMP-PBMCsec-production: leukocyte cones are obtained as blood-donation by-product, PBMCs are isolated by Ficoll-centrifugation, cells are irradiated with 60Gy and incubated for 24h. Superantatants are filtrated and lyophilized, and off-the-shelf PBMCsec is stored at -20° until use.
B) Table of antibodies used. IF = immunofluorescence, WB = western blot

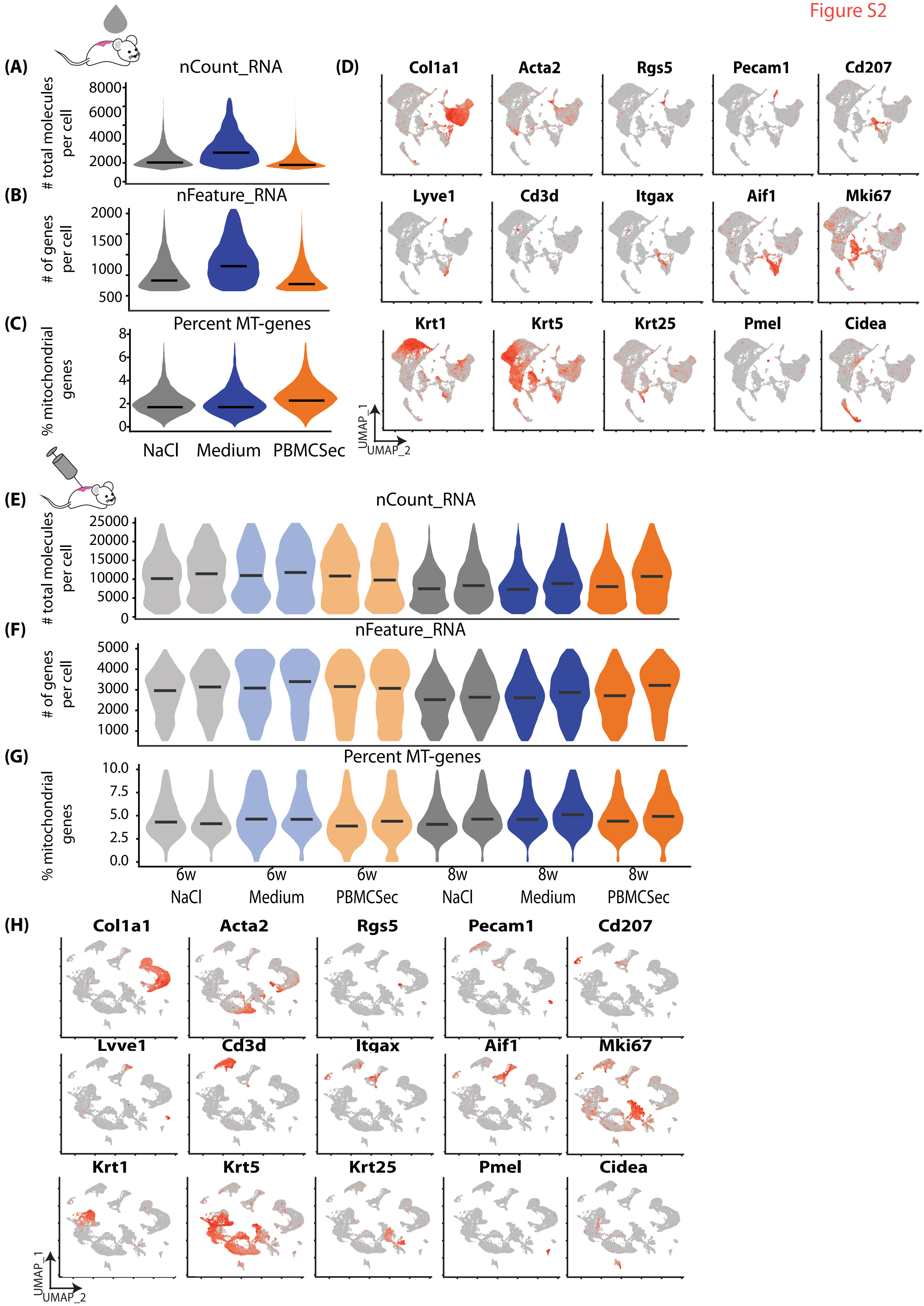
(A)
nCount_RNA
### total molecules per cell
8000
6000
4000
2000
0
(B)
nFeature_RNA
### of genes per cell
2000
1500
1000
500
(C)
Percent MT-genes
% mitochondrial genes
8
6
4
2
0
NaCl
Medium
PBMCSec
(D)
Col1a1
Acta2
Rgs5
Pecam1
Cd207
Lyve1
Cd3d
Itgax
Aif1
Mki67
Krt1
Krt5
Krt25
Pmel
Cidea
UMAP_1
UMAP_2
(E)
nCount_RNA
### total molecules per cell
25000
20000
15000
10000
5000
0
(F)
nFeature_RNA
### of genes per cell
5000
4000
3000
2000
1000
(G)
Percent MT-genes
% mitochondrial genes
10.0
7.5
5.0
2.5
0.0
6w NaCl
6w Medium
6w PBMCSec
8w NaCl
8w Medium
8w PBMCSec
(H)
Col1a1
Acta2
Rgs5
Pecam1
Cd207
Lyve1
Cd3d
Itgax
Aif1
Mki67
Krt1
Krt5
Krt25
Pmel
Cidea
UMAP_1
UMAP_2

**Figure S2: Quality control of mouse scRNAseq**

A-C, E-G) Violin plots of quality control parameters of the 'topical' and 'inject' dataset. A, E) total molecules per cell B, F) (gene count per cell and C, G) mitochondrial gene content. D, H) Feature Plots of cluster markers for cluster identification in the 'topical' and 'inject' dataset: *Col1a1* (collagen I alpha 1) for fibroblasts, *Acta2* (smooth muscle actin) for smooth muscle cells and myofibroblasts, *Rgs5* (Regulator Of G Protein Signaling 5) for pericytes, *Pecam* (Platelet And Endothelial Cell Adhesion Molecule 1) for endothelial cells, *Lyve1* (Lymphatic Vessel Endothelial Hyaluronan Receptor 1) for lymphatic endothelial cells, *Cd207* (Langerin) for Langerhans cells, *Cd3d* (cluster of differentiation 3D) for T-cells, *Itgax* for dendritic cells, *Aif1* (allograft inflammatory factor 1) for macrophages, *Mki67* (Marker Of Proliferation Ki-67) for proliferating cells, *Krt1* (Keratin1) for spinous and granular keratinocytes (KCs), *Krt5* (Keratin 5) for basal KCs, *Krt25* (Keratin 25) for hair follicles, *Pmel* (Premelanosome Protein) for melanocytes, *Cidea* (Cell Death Inducing DFFA Like Effector A) for adipocytes. Vertical lines in violin plots represent maximum expression, shape of each violin represents all results, and width of each violin represents frequency of cells at the respective expression level. In feature plots, normalized log expression of the respective gene is mapped onto the UMAP-plot. Color intensity indicates level of gene expressions. UMAP, uniform manifold approximation and projection.

Figure S3

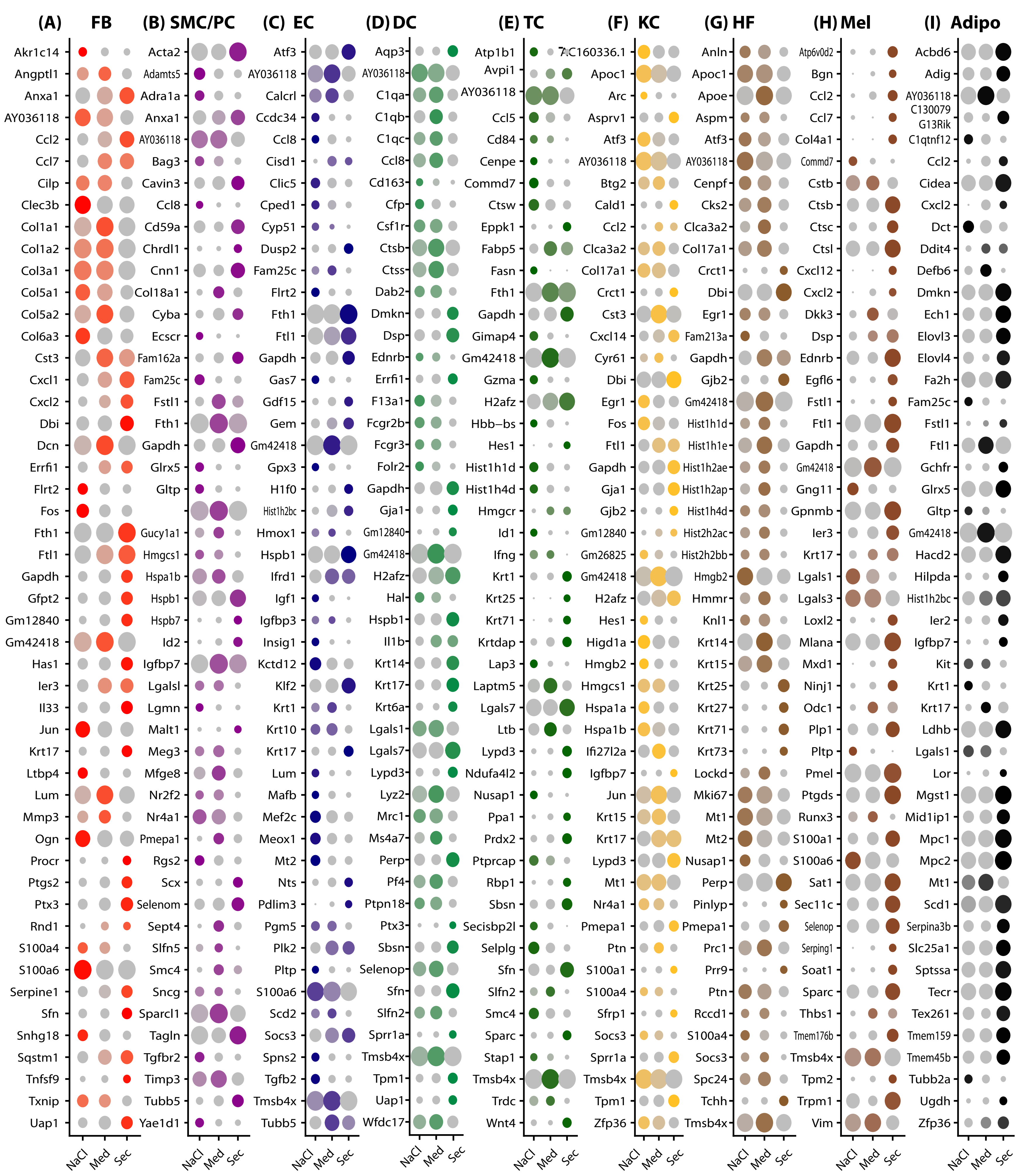
(A) FB
Akr1c14
Angptl1
Anxa1
AY036118
Ccl2
Ccl7
Cilp
Clec3b
Col1a1
Col1a2
Col3a1
Col5a1
Col5a2
Col6a3
Cst3
Cxcl1
Cxcl2
Dbi
Dcn
Errfi1
Flrt2
Fos
Fth1
Ftl1
Gapdh
Gfpt2
Gm12840
Gm42418
Has1
Ier3
Il33
Jun
Krt17
Ltbp4
Lum
Mmp3
Ogn
Procr
Ptgs2
Ptx3
Rnd1
S100a4
S100a6
Serpine1
Sfn
Snhg18
Sqstm1
Tnfsf9
Txnip
Uap1
(B) SMC/PC
Acta2
Adamts5
Adra1a
Anxa1
AY036118
Bag3
Cavin3
Ccl8
Cd59a
Chrdl1
Cnn1
Col18a1
Cyba
Ecscr
Fam162a
Fam25c
Fstl1
Fth1
Gapdh
Glrx5
Gltp
Gucy1a1
Hmgcs1
Hspa1b
Hspb1
Hspb7
Id2
Igfbp7
Lgalsl
Lgmn
Malt1
Meg3
Mfge8
Nr2f2
Nr4a1
Pmepa1
Rgs2
Scx
Selenom
Sept4
Slfn5
Smc4
Sncg
Sparcl1
Tagln
Tgfbr2
Timp3
Tubb5
Yae1d1
(C) EC
Atf3
AY036118
Calcrl
Ccdc34
Ccl8
Cisd1
Clic5
Cped1
Cyp51
Dusp2
Fam25c
Flrt2
Fth1
Ftl1
Gapdh
Gas7
Gdf15
Gem
Gm42418
Gpx3
H1f0
Hist1h2bc
Hmox1
Hspb1
Ifrd1
Igf1
Igfbp3
Insig1
Kctd12
Klf2
Krt1
Krt10
Krt17
Lum
Mafb
Mef2c
Meox1
Mt2
Nts
Pdlim3
Pgm5
Plk2
Pltp
S100a6
Scd2
Socs3
Spns2
Tgfb2
Tmsb4x
Tubb5
(D) DC
Aqp3
AY036118
C1qa
C1qb
C1qc
Ccl8
Cd163
Cfp
Csf1r
Ctsb
Ctss
Dab2
Dmkn
Dsp
Ednrb
Errfi1
F13a1
Fcgr2b
Fcgr3
Folr2
Gapdh
Gja1
Gm12840
Gm42418
H2afz
Hal
Hspb1
Il1b
Krt14
Krt17
Krt6a
Lgals1
Lgals7
Lypd3
Lyz2
Mrc1
Ms4a7
Perp
Pf4
Ptpn18
Ptx3
Sbsn
Selenop
Sfn
Slfn2
Sprr1a
Tmsb4x
Tpm1
Uap1
Wfdc17
(E) TC
Atp1b1
Avpi1
AY036118
Ccl5
Cd84
Cenpe
Commd7
Ctsw
Eppk1
Fabp5
Fasn
Fth1
Gapdh
Gimap4
Gm42418
Gzma
H2afz
Hbb−bs
Hes1
Hist1h1d
Hist1h4d
Hmgcr
Id1
Ifng
Krt1
Krt25
Krt71
Krtdap
Lap3
Laptm5
Lgals7
Ltb
Lypd3
Ndufa4l2
Nusap1
Ppa1
Prdx2
Ptprcap
Rbp1
Sbsn
Secisbp2l
Selplg
Sfn
Slfn2
Smc4
Sparc
Stap1
Tmsb4x
Trdc
Wnt4
(F) KC
AC160336.1
Apoc1
Arc
Asprv1
Atf3
AY036118
Btg2
Cald1
Ccl2
Clca3a2
Col17a1
Crct1
Cst3
Cxcl14
Cyr61
Dbi
Egr1
Fos
Ftl1
Gapdh
Gja1
Gjb2
Gm12840
Gm26825
Gm42418
H2afz
Hes1
Higd1a
Hmgb2
Hmgcs1
Hspa1a
Hspa1b
Ifi27l2a
Igfbp7
Jun
Krt15
Krt17
Lypd3
Mt1
Nr4a1
Pmepa1
Ptn
S100a1
S100a4
Sfrp1
Socs3
Sprr1a
Tmsb4x
Tpm1
Zfp36
(G) HF
Anln
Apoc1
Apoe
Aspm
Atf3
AY036118
Cenpf
Cks2
Clca3a2
Col17a1
Crct1
Dbi
Egr1
Fam213a
Gapdh
Gjb2
Gm42418
Hist1h1d
Hist1h1e
Hist1h2ae
Hist1h2ap
Hist1h4d
Hist2h2ac
Hist2h2bb
Hmgb2
Hmmr
Knl1
Krt14
Krt15
Krt25
Krt27
Krt71
Krt73
Lockd
Mki67
Mt1
Mt2
Nusap1
Perp
Pinlyp
Pmepa1
Prc1
Prr9
Ptn
Rccd1
S100a4
Socs3
Spc24
Tchh
Tmsb4x
(H) Mel
Atp6v0d2
Bgn
Ccl2
Ccl7
Col4a1
Commd7
Cstb
Ctsb
Ctsc
Ctsl
Cxcl12
Cxcl2
Dkk3
Dsp
Ednrb
Egfl6
Fstl1
Ftl1
Gapdh
Gm42418
Gng11
Gpnmb
Ier3
Krt17
Lgals1
Lgals3
Loxl2
Mlana
Mxd1
Ninj1
Odc1
Plp1
Pltp
Pmel
Ptgds
Runx3
S100a1
S100a6
Sat1
Sec11c
Selenop
Serping1
Soat1
Sparc
Thbs1
Tmem176b
Tmsb4x
Tpm2
Trpm1
Vim
(I) Adipo
Acbd6
Adig
AY036118
C130079
G13Rik
C1qtnf12
Ccl2
Cidea
Cxcl2
Dct
Ddit4
Defb6
Dmkn
Ech1
Elovl3
Elovl4
Fa2h
Fam25c
Fstl1
Ftl1
Gchfr
Glrx5
Gltp
Gm42418
Hacd2
Hilpda
Hist1h2bc
Ier2
Igfbp7
Kit
Krt1
Krt17
Ldhb
Lgals1
Lor
Mgst1
Mid1ip1
Mpc1
Mpc2
Mt1
Scd1
Serpina3b
Slc25a1
Sptssa
Tecr
Tex261
Tmem159
Tmem45b
Tubb2a
Ugdh
Zfp36
NaCl
Med
Sec

**Figure S3**: **Top 50 regulated genes per cell group in 'Topical' mouse scars**

In A) fibroblasts (FBs, red circles), B) smooth muscle cells and pericytes (SMC/PCs, purple), C) endothelial cells (ECs, blue), D) T-cells (dark green), E) dendritic cells and macrophages (DCs, light green), and keratinocytes (KCs, yellow), hair follicles (HF, beige), melanocytes (Mel, brown), adipocytes (Adipo, grey); differentially expressed genes (DEGs) were calculated comparing 'PBMCsec' mouse scars to 'NaCl'-mouse scars, using Wilcoxon rank sum test, including genes with average logarithmic fold change (avglogFC) of > 0.1 or < -0.1 and Bonferroni-adjusted p-value <0.05. For each cellgroup, top 50 DEGs according to lowest adjusted p-value are displayed, split by treatment. Dot size represents percent of cells expressing the respective gene, color correlates with average expression.

Figure S4

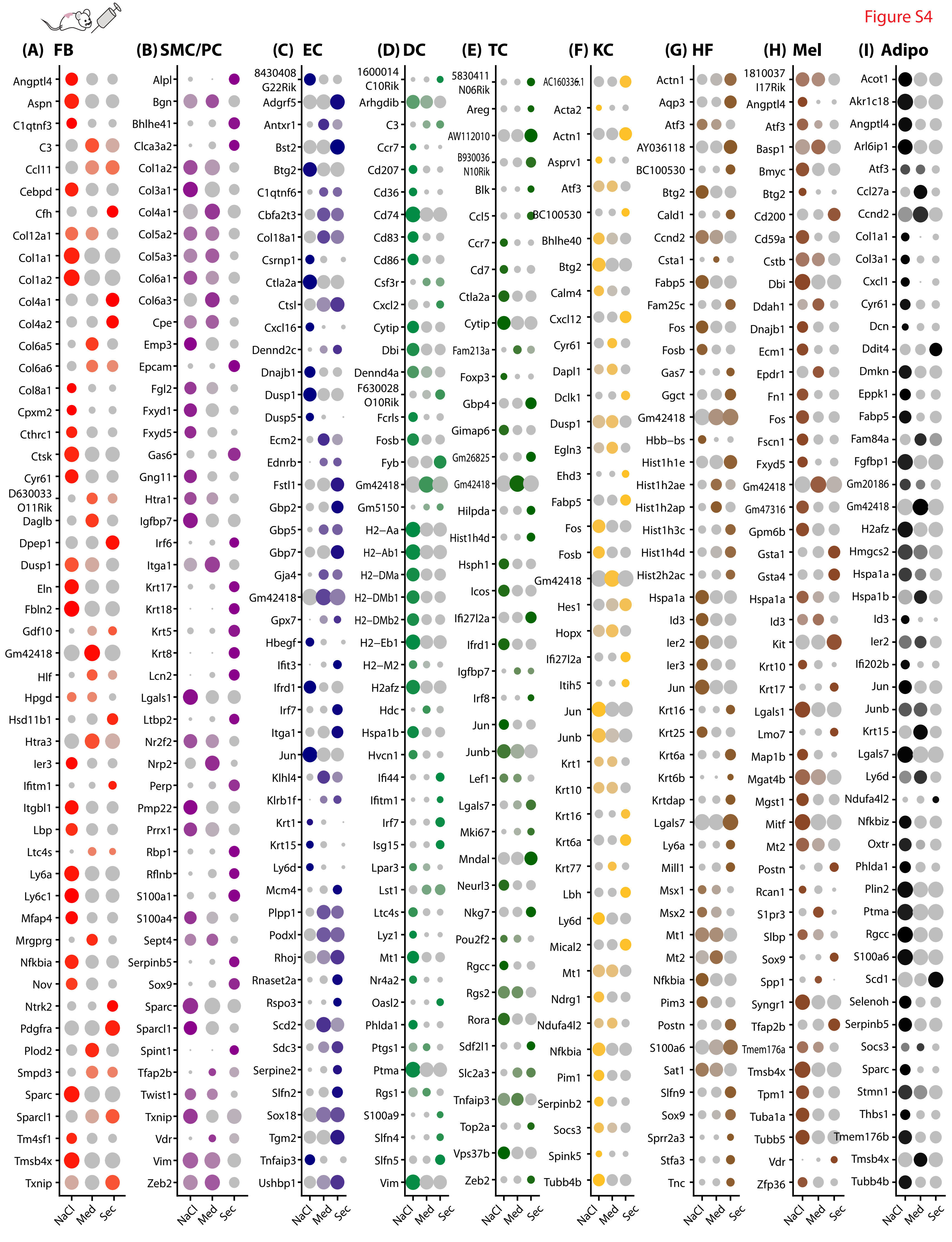

**Figure S4: Top 50 regulated genes per cell group in 6 weeks 'Inject' mouse scars**

In A) fibroblasts (FBs, red circles), B) smooth muscle cells and pericytes (SMC/PCs, purple), C) endothelial cells (ECs, blue), D) T-cells (dark green), E) dendritic cells and macrophages (DCs, light green), and keratinocytes (KCs, yellow), hair follicles (HF, beige), melanocytes (Mel, brown), adipocytes (Adipo, grey); differentially expressed genes (DEGs) were calculated comparing 'PBMCsec' mouse scars to 'NaCl'-mouse scars, using Wilcoxon rank sum test, including genes with average logarithmic fold change (avglogFC) of > 0.1 or < -0.1 and Bonferroni-adjusted p-value <0.05. For each cellgroup, top 50 DEGs according to lowest adjusted p-value are displayed, split by treatment and timepoint. Dot size represents percent of cells expressing the respective gene, color correlates with average expression.

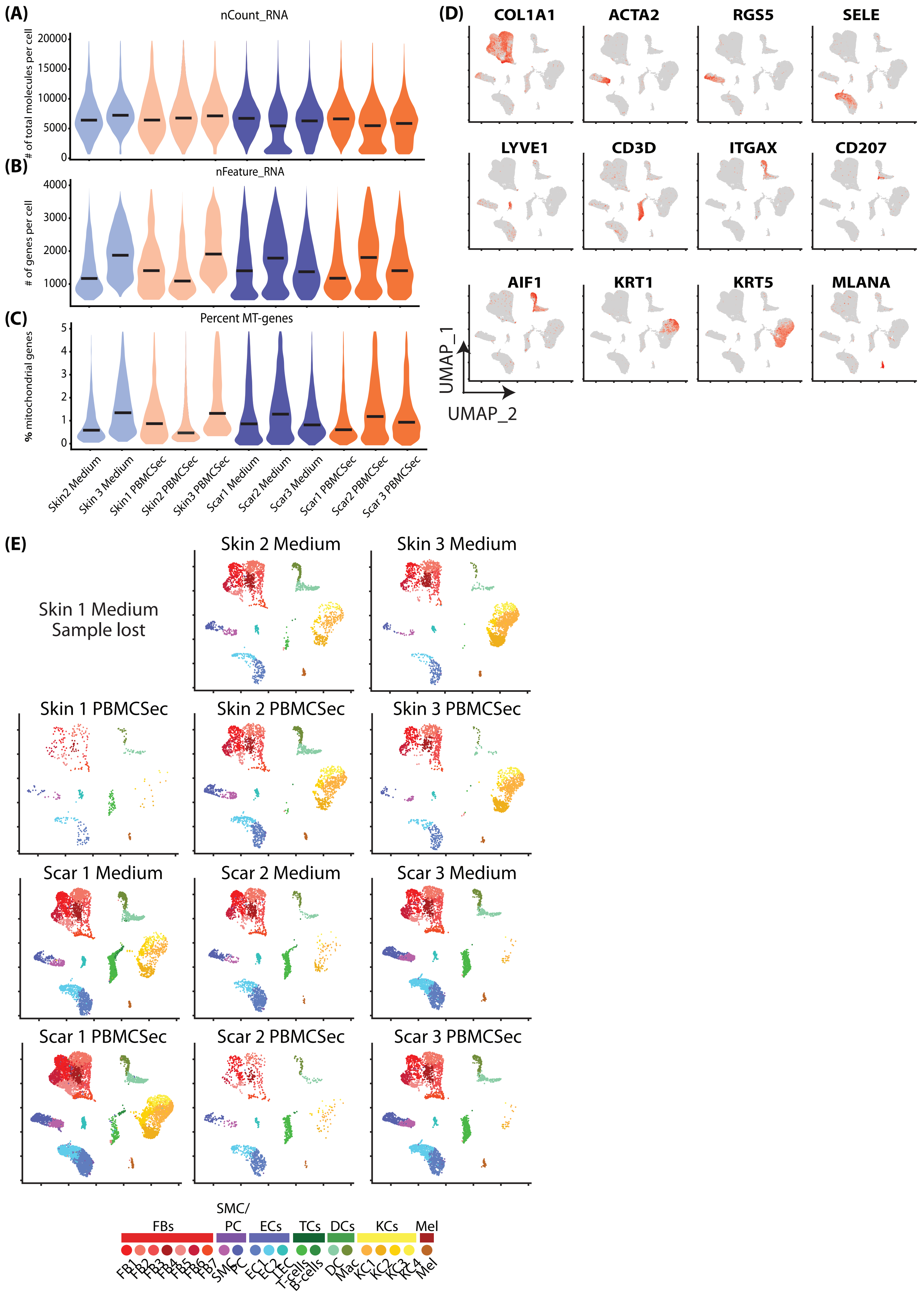
(A)
nCount_RNA
### of total molecules per cell
20000
15000
10000
5000
0
(B)
nFeature_RNA
### of genes per cell
4000
3000
2000
1000
(C)
Percent MT-genes
% mitochondrial genes
5
4
3
2
1
0
Skin2 Medium
Skin 3 Medium
Skin1 PBMCSec
Skin2 PBMCSec
Skin3 PBMCSec
Scar1 Medium
Scar2 Medium
Scar3 Medium
Scar1 PBMCSec
Scar2 PBMCSec
Scar 3 PBMCSec
(D)
COL1A1
ACTA2
RGS5
SELE
LYVE1
CD3D
ITGAX
CD207
AIF1
KRT1
KRT5
MLANA
UMAP_1
UMAP_2
(E)
Skin 1 Medium
Sample lost
Skin 2 Medium
Skin 3 Medium
Skin 1 PBMCSec
Skin 2 PBMCSec
Skin 3 PBMCSec
Scar 1 Medium
Scar 2 Medium
Scar 3 Medium
Scar 1 PBMCSec
Scar 2 PBMCSec
Scar 3 PBMCSec
FBs
SMC/ PC
ECs
TCs
DCs
KCs
Mel
FB1
FB2
FB3
FB4
FB5
FB6
FB7
SMC
PC
EC1
EC2
LEC
T-cells
B-cells
DC
Mac
KC1
KC2
KC3
KC4
Mel

**Figure S5: Quality control of human scRNAseq**

A-C) Violin plots of quality control parameters of the 'human' dataset. A) total molecules per cell B) (gene count per cell and C) mitochondrial gene content. D) Feature Plots of cluster markers for cluster identification in the 'topical' and 'inject' dataset: *COL1A1* (collagen I alpha 1) for fibroblasts, *ACTA2* (smooth muscle actin) for smooth muscle cells and myofibroblasts, *RGS5* (Regulator Of G Protein Signaling 5) for pericytes, *SELE* (Selectin E) for endothelial cells, *LYVE1* (Lymphatic Vessel Endothelial Hyaluronan Receptor 1) for lymphatic endothelial cells, *Cd3d* (cluster of differentiation 3D) for T-cells, *ITGAX* for dendritic cells, *CD207* (Langerin) for Langerhans cells, *AIF1* (allograft inflammatory factor 1) for macrophages, *KRT1* (Keratin1) for spinous and granular keratinocytes (KCs), *KRT5* (Keratin 5) for basal KCs, *MLANA* (Melan-A) for melanocytes. E) UMAP-clustering in the 'human' dataset, split by samples. Sample "Skin 1 medium" was lost due to technical difficulties during preparation. Vertical lines in violin plots represent maximum expression, shape of each violin represents all results, and width of each violin represents frequency of cells at the respective expression level. In feature plots, normalized log expression of the respective gene is mapped onto the UMAP-plot. Color intensity indicates level of gene expressions. UMAP, uniform manifold approximation and projection

Figure S6 skin

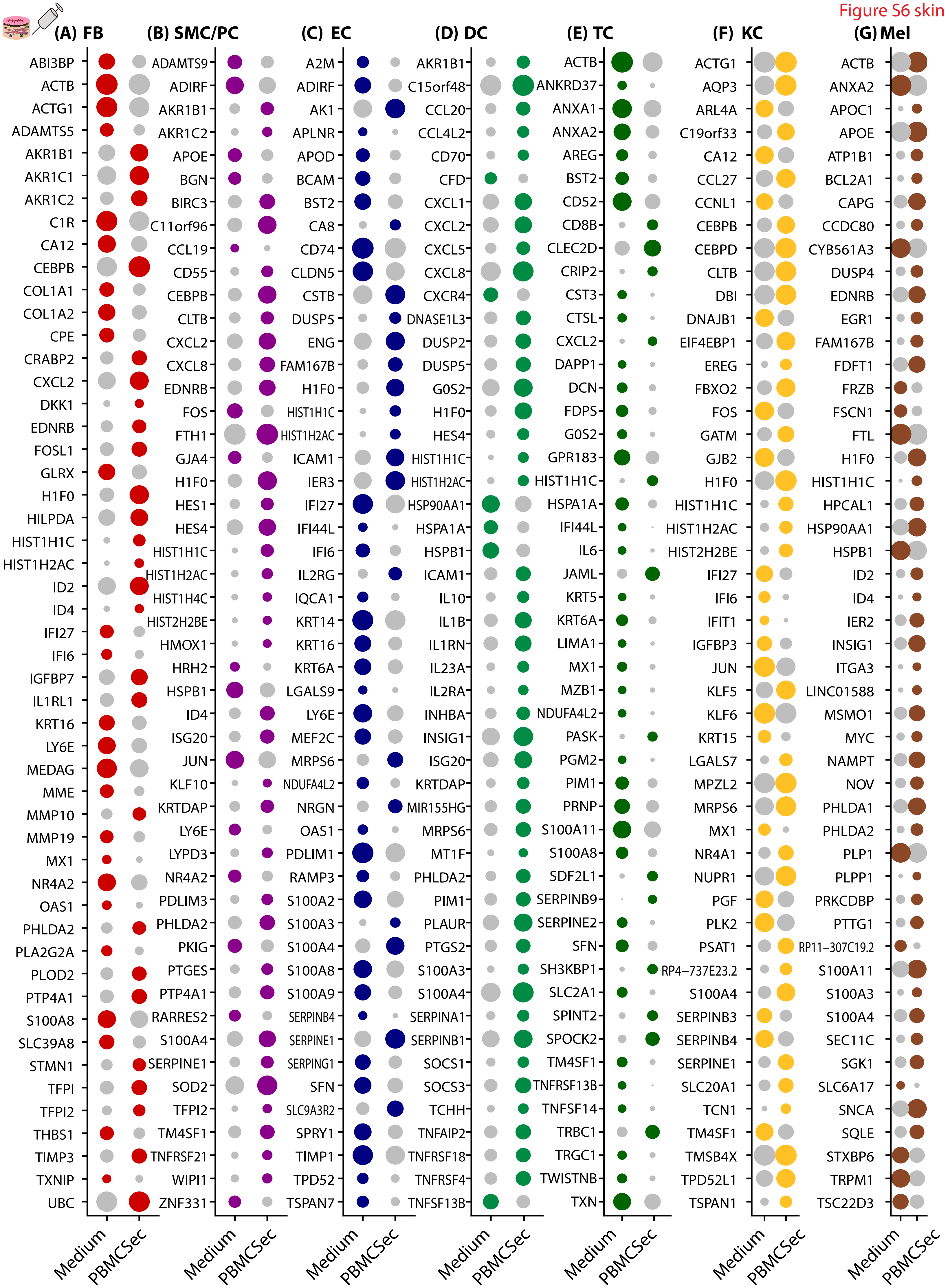

**Figure S6: Top 50 regulated genes per cell group in human skin**

In A) fibroblasts (FBs, red circles), B) smooth muscle cells and pericytes (SMC/PCs, purple), C) endothelial cells (ECs, blue), D) T-cells (dark green), E) dendritic cells and macrophages (DCs, light green), and keratinocytes (KCs, yellow), melanocytes (Mel, brown); differentially expressed genes (DEGs) were calculated comparing 'PBMCsec' skin biopsies to 'Medium' skin biopsies, using Wilcoxon rank sum test, including genes with average logarithmic fold change (avglogFC) of > 0.1 or < -0.1 and Bonferroni-adjusted p-value <0.05. For each cellgroup, top 50 DEGs according to lowest adjusted p-value are displayed, split by treatment. Dot size represents percent of cells expressing the respective gene, color correlates with average expression.

Figure S7 scar

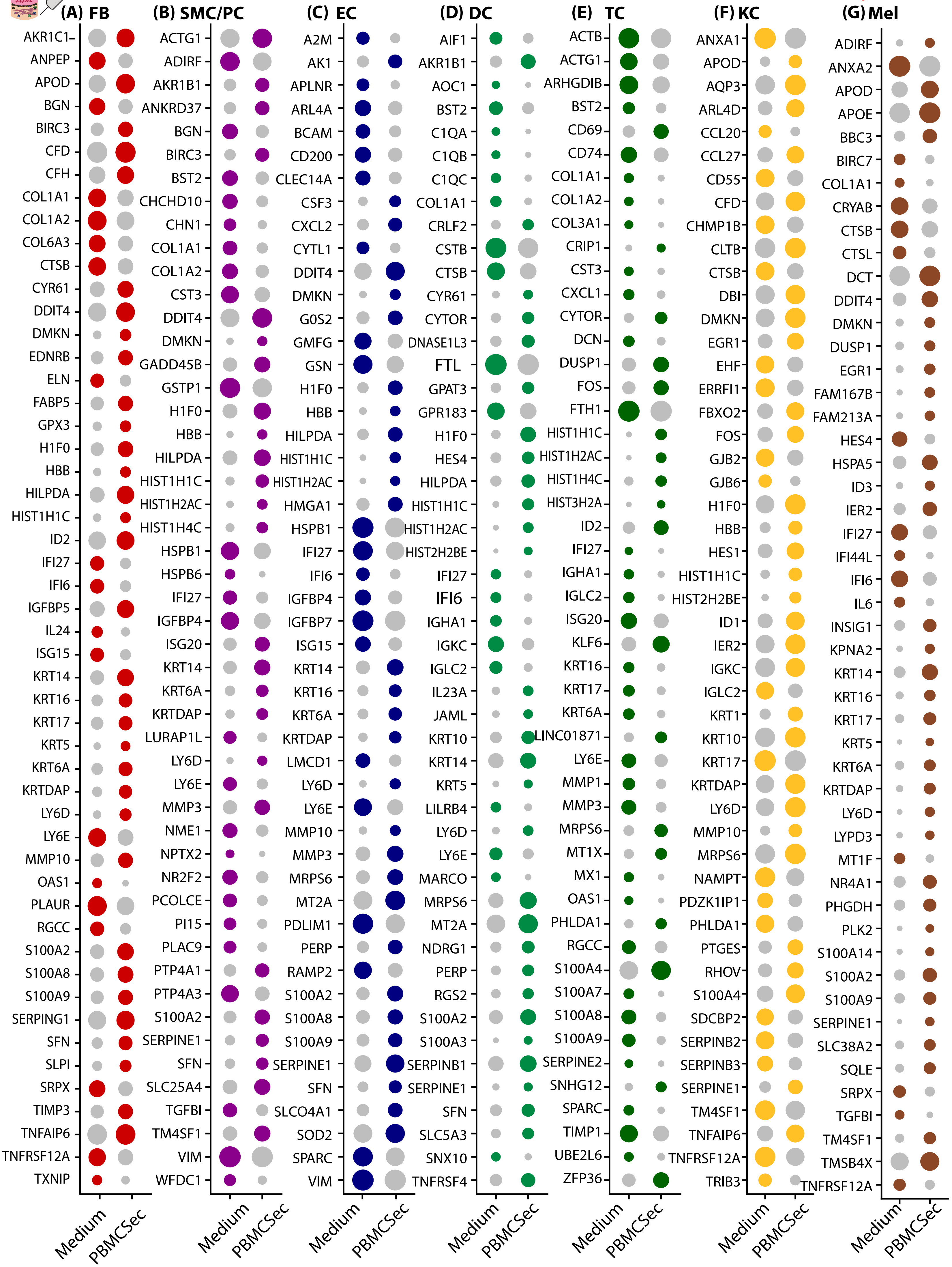

**Figure S7: Top 50 regulated genes per cell group in human scar**

In A) fibroblasts (FBs, red circles), B) smooth muscle cells and pericytes (SMC/PCs, purple), C) endothelial cells (ECs, blue), D) T-cells (dark green), E) dendritic cells and macrophages (DCs, light green), and keratinocytes (KCs, yellow), melanocytes (Mel, brown); differentially expressed genes (DEGs) were calculated comparing ‘PBMCsec’ scar biopsies to ‘Medium’ scar biopsies, using Wilcoxon rank sum test, including genes with average logarithmic fold change (avglogFC) of > 0.1 or < -0.1 and Bonferroni-adjusted p-value <0.05. For each cellgroup, top 50 DEGs according to lowest adjusted p-value are displayed, split by treatment. Dot size represents percent of cells expressing the respective gene, color correlates with average expression.

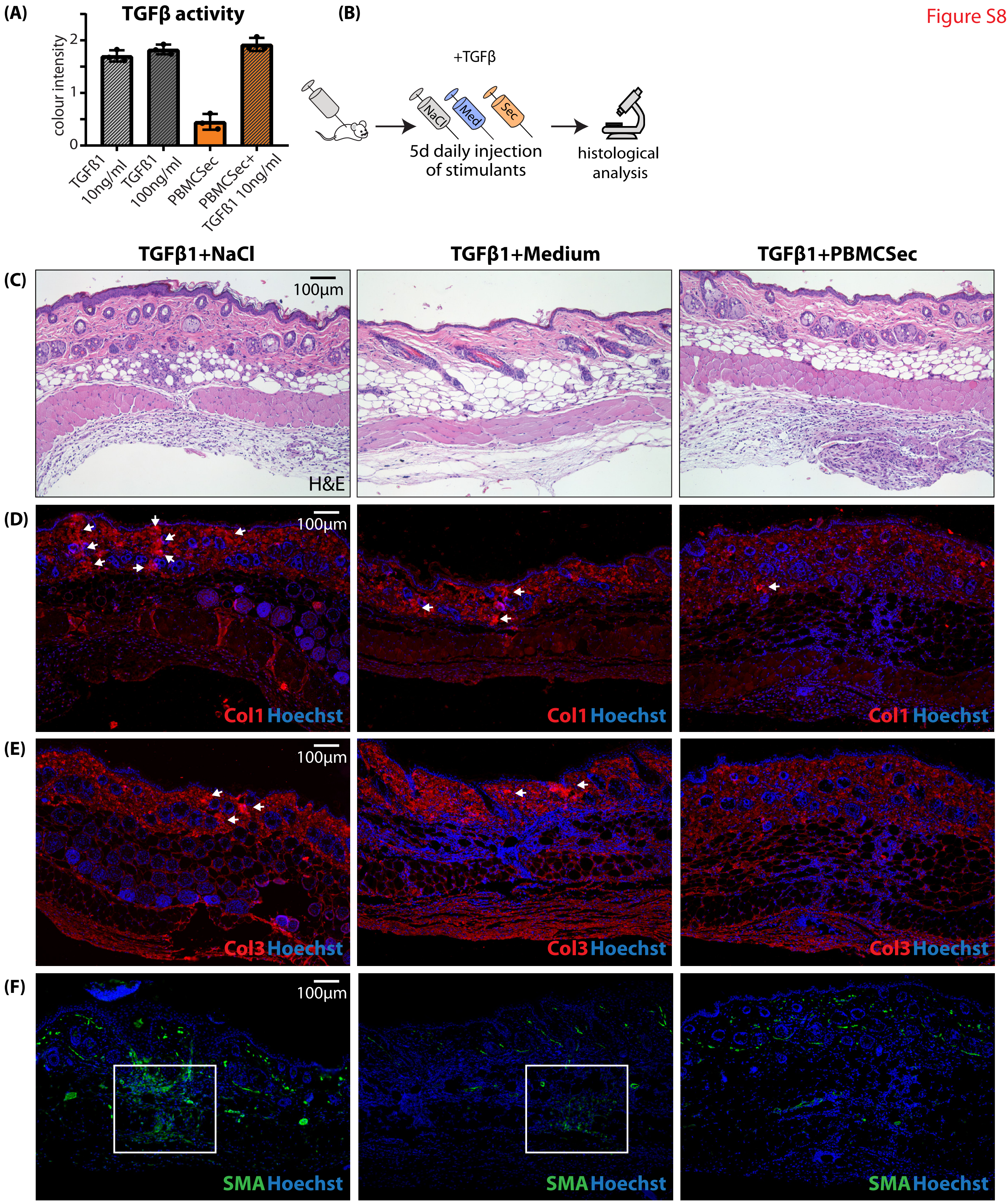
(A)
TGFβ activity
colour intensity
0
1
2
TGFβ1 10ng/ml
TGFβ1 100ng/ml
PBMCSec
PBMCSec+ TGFβ1 10ng/ml
(B)
+TGFβ
NaCl
Med
Sec
5d daily injection of stimulants
histological analysis
TGFβ1+NaCl
TGFβ1+Medium
TGFβ1+PBMCSec
(C)
100µm
H&E
(D)
100µm
Col1 Hoechst
Col1 Hoechst
Col1 Hoechst
(E)
100µm
Col3 Hoechst
Col3 Hoechst
Col3 Hoechst
(F)
100µm
SMA Hoechst
SMA Hoechst
SMA Hoechst

**Figure S8: The interaction of TGFβ and PBMCsec *in vivo***

A) In vitro TGFβ-activity assay. HEK-cells colorimetrically detecting SMAD2/3 TGFβ-activity were incubated with recombinant TGFβ1, with PBMCsec alone, and with TGFβ1 and PBMCsec combined. Color intensity correlates with TGFβ1-signaling activity. B) Workflow of mice intradermally injected with TGFβ1 and treatmens. Mice were intradermally injected with 800ng TGFβ1 dissolved 100µl in NaCl 0,9%, Medium or PBMCsec on five consecutive days and sacrificed on the $6^{th}$ day. C) H&E staining of resulting “scars” of the injected area. Immunofluorescence stainings of D) collagen 1, E) collagen 3, F) smooth muscle actin in TGFβ1-injected mouse skin. Arrows indicate areas of dense matrix deposition.

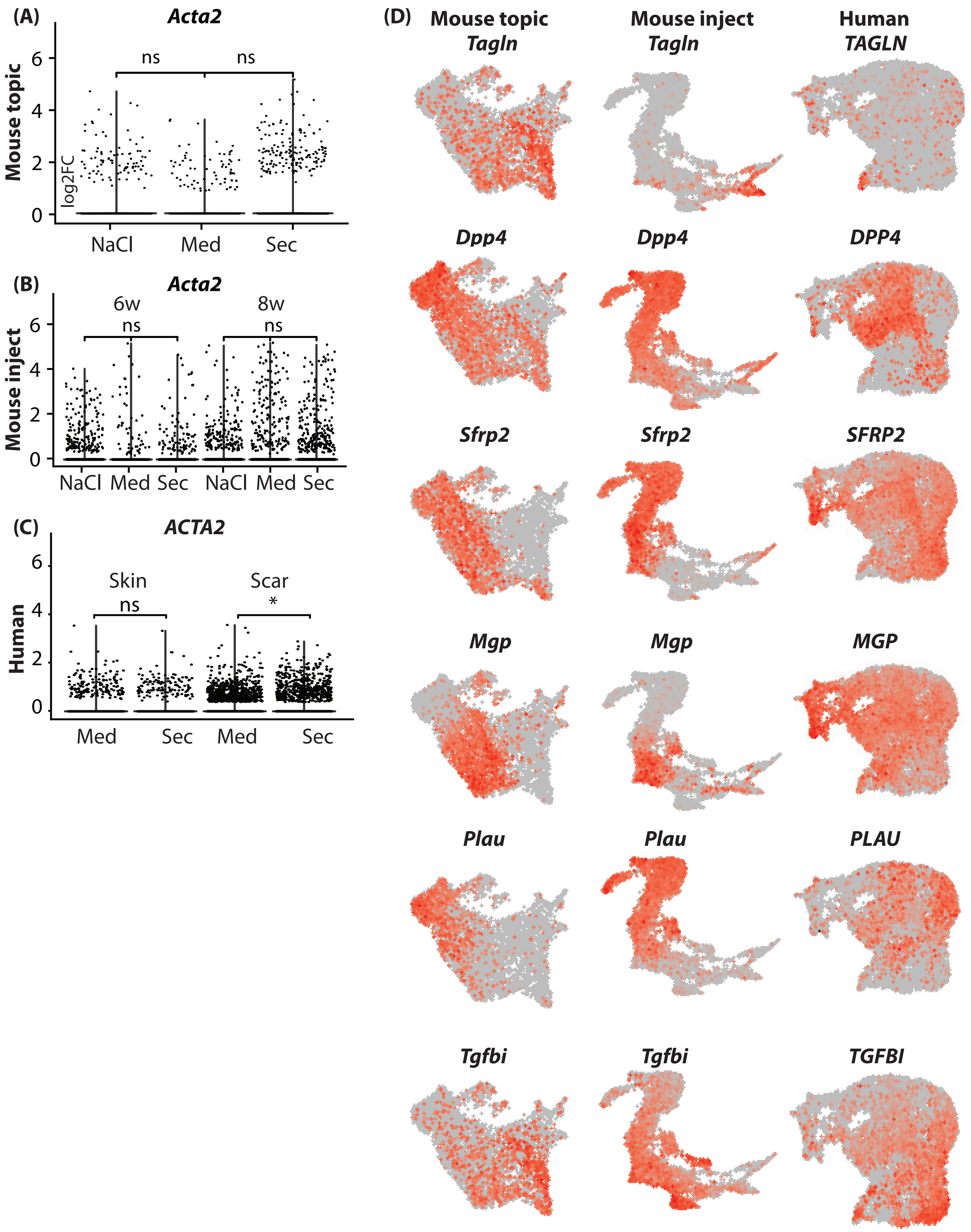
(A)
Acta2
Mouse topic
log2FC
ns
ns
NaCl
Med
Sec
(B)
Acta2
Mouse inject
6w
ns
8w
ns
NaCl
Med
Sec
NaCl
Med
Sec
(C)
ACTA2
Human
Skin
ns
Scar
*
Med
Sec
Med
Sec
(D)
Mouse topic
Mouse inject
Human
Tagln
Tagln
TAGLN
Dpp4
Dpp4
DPP4
Sfrp2
Sfrp2
SFRP2
Mgp
Mgp
MGP
Plau
Plau
PLAU
Tgfbi
Tgfbi
TGFBI

**Figure S9: Subcluster analysis of FB populations**

A-C) Violin plots of *Acta2/ACTA2* in mouse and human datasets, split by treatments. D) Feature plots of FB subcluster markers in FB subclusters of mouse and human datasets. *Tagln/TAGLN*, Transgelin; *Dpp4/DPP4*, dipeptidyl-peptidase 4; *Sfrp2/SFRP2,* Secreted Frizzled Related Protein 2; *Mgp/MGP,* Matrix Gla Protein; *Plau/PLAU*, urokinase; *Tgfbi/TGFBI*, Transforming Growth Factor Beta Induced. In violin plots, dots represent individual cells, *y*-axis represents log2 fold change of the normalized genes and log-transformed single-cell expression. Vertical lines in violin plots represent maximum expression, shape of each violin represents all results, and width of each violin represents frequency of cells at the respective expression level. In feature plots, normalized log expression of the respective gene is mapped onto the UMAP-plot. Color intensity indicates level of gene expressions. UMAP, uniform manifold approximation and projection. A two-sided Wilcoxon-signed rank test was used in R. NS $p > 0.05$, *$p < 0.05$, **$p < 0.01$, ***$p < 0.001$.
